# Legacy Effects of Land-Use Land-Cover Change on Species Distribution Dynamics: Evidence from a landscape of diverse biogeographic crossroads

**DOI:** 10.64898/2026.06.01.729351

**Authors:** Shrutarshi Paul, Vivianna Borzym, Heather Prestridge, Wenzhe Jiao, Mary Katherine Gonder

**Author notes:** Corresponding authors: Shrutarshi Paul, Mary Katherine Gonder.

## Abstract

Land-use and land-cover (LULC) change is a dominant driver of biodiversity loss, yet its long-term role in restructuring species distribution remains poorly understood due to limited historical baselines and the scarcity of long-term analyses across multiple ecological transition zones. Here, we investigate nearly two centuries of land-use change across a biogeographically diverse region to evaluate how landscape transformation has reshaped vertebrate distribution range dynamics, and whether ecoregional context, habitat specialization, and taxonomic identity better explain these patterns than recent landscape change alone. Using Texas, a biogeographic crossroads with diverse ecoregional zones and a long history of intensive land-use, we quantify distributional changes in 100 native vertebrate species (mammals, birds, reptiles, and amphibians), using Vernon Bailey’s late nineteenth-century surveys as a historical benchmark. We integrated multi-temporal LULC datasets with spatial and multivariate analyses to assess how habitat change, species traits, and regional context relate to patterns of range expansion, contraction, and redistribution. We show that Texas has undergone non-linear landscape transformations characterized by early agricultural expansion followed by cropland abandonment, shrubland encroachment, and rapid urbanization. Range contractions were most pronounced among amphibians, reptiles, and habitat specialists, whereas generalist species and many birds exhibited greater range stability or expansion. Across taxa, primary habitat, vertebrate class, and ecoregional context emerged as the strongest predictors of distributional change, underscoring the importance of historical and environmental context over simple measures of habitat loss and fragmentation. These results indicate that while long-term LULC change establishes the underlying landscape template, species responses are structured by ecological context, including habitat affinity, taxonomic traits, and regional environmental gradients. By linking historical landscape trajectories to contemporary biodiversity patterns, this study provides a transferable framework for investigating how land-use legacies influence species distributions and community reassembly in heterogeneous human-dominated landscapes, highlighting the importance of adaptive capacity and connectivity for conservation planning.

## 1. Introduction

Land-use and land-cover (LULC) change is one of the most pervasive drivers of global biodiversity loss, restructuring ecosystems at spatial and temporal scales comparable to, and often interacting with, climate change (Diaz et al., 2019; Jung et al., 2020; Wang et al., 2022). Agricultural expansion, rangeland modification, urbanization, infrastructural development, and resource extraction have transformed large portions of Earth’s surface into heterogeneous mosaics, altering habitat availability, connectivity, and disturbance regimes worldwide (Kuykendall & Gregory, 2011; Mullu, 2016; Ren et al., 2022; Surasinghe & Baldwin, 2014). Rather than affecting species independently, these transformations drive shifts in species distributions through range contractions, expansions, and reorganization, with implications for community restructuring, ecosystem functioning, evolutionary trajectories, and resilience to future environmental change (Murphy et al., 2025; Newbold et al., 2019; Tewksbury et al., 2002).

From a community-assembly perspective, land-use change acts as a form of ecological filtering, favoring species with traits that enable persistence in modified landscapes, such as broad habitat tolerance, high dispersal capacity, and behavioral flexibility, while constraining habitat specialists and less mobile taxa. As a result, anthropogenic landscapes increasingly support novel assemblages composed of persistent generalists and declining specialists, reflecting directional shifts in species composition (Clavel et al., 2011; Newbold et al., 2019).

Globally, responses to landscape transformation are highly uneven (Daskalova et al., 2020). About one-third of vertebrates are experiencing range contractions globally (Brooks et al., 2002; Ceballos et al., 2017; Jung et al., 2020; Pimm et al., 2014; Vargas-Jaimes et al., 2021; WWF, 2018), a pattern particularly pronounced among habitat specialists, whereas generalists and highly mobile species may persist or even expand their ranges. These effects are amplified in human-modified landscapes with strong environmental gradients, biogeographic transitions or biogeographic crossroads characterized by complex interplay of ecological zones and species assemblages, in which species distributions are already constrained by steep climatic and ecological boundaries (Gaston, 2009; Jung et al., 2020). In such systems, land-use change can accelerate range shifts and reorganize species distributions, making them especially sensitive to sustained anthropogenic pressures (Newbold et al., 2019). Understanding these dynamics requires a long-term perspective and is increasingly recognized as vital for guiding restoration priorities, identifying conservation hotspots, and forecasting future ecological trajectories.

However, most studies of biodiversity responses to land-use change rely on contemporary or short-term datasets, limiting our ability to distinguish directional, human-driven change from natural variability and to quantify legacy effects of past land-use. Integrating historical baselines with contemporary data provides a critical opportunity to examine how species distribution range changes over extended time scales and to identify the mechanisms driving differential species responses (Fukasawa & Akasaka, 2019; Surasinghe & Baldwin, 2014). Regions that combine pronounced environmental heterogeneity with well-documented histories of land-use change offer particularly powerful natural laboratories for testing these ideas (De Frenne et al., 2013; García-Vega & Newbold, 2020; Kéfi et al., 2024).

The state of Texas exemplifies such a system, widely regarded as a biogeographic crossroads where eastern forests transition into western deserts, east–west precipitation gradients are steep, and proximity to the tropics creates complex environmental heterogeneity and diverse species assemblages (TPWD, 2002; Webb, 1950). The state harbors some of the highest levels of biodiversity and endemism in the United States and encompasses ten distinct ecoregions with divergent ecological conditions (Schmandt et al., 2011). Prior to Euro-American settlement, many of these ecosystems were shaped by long-standing indigenous land stewardship, including the use of frequent, low-intensity fire that maintained grasslands, limited woody encroachment, and contributed to heterogeneous habitat mosaics (Pyne, 1997; Roos et al., 2018; Swetnam et al., 2016). The transition to industrial land-use, therefore, represents not the onset of human influence but a fundamental shift in its intensity and ecological consequences. Since the mid-nineteenth century, Texas has undergone extensive and spatially heterogeneous transformations, including agricultural expansion, rangeland alteration, and rapid urbanization (Johnston, 2014; Lombardi et al., 2020; Wilcox et al., 2012). Nevertheless, the long-term consequences of these transformations for species distributions remain incompletely understood. Texas offers a rare combination of historical biological surveys, LULC reconstructions, and contemporary biodiversity datasets. Early faunal surveys conducted by Vernon Bailey through the Biological Survey of Texas in the late nineteenth century provide a pre-modern baseline for species distributions (Bailey, 1905), while recent advances in spatial data allow reconstructions of landscape change from the mid-nineteenth century to the present and recent biological surveys. Together, these data provide a unique opportunity to examine how nearly two centuries of landscape transformation have shaped patterns of vertebrate distribution with implications for enhancing understanding of principles of community reassembly.

Critically, more than 95% of Texas land is privately owned (Kreuter et al., 2017), creating a distinctive socioecological dimension in which land management decisions across decentralized stakeholders directly shape habitat availability and biodiversity patterns. Policy mechanisms such as wildlife management valuation and tax-incentivized conservation further influence these dynamics by shaping land-use practices across a fragmented ownership mosaic (Benavidez et al., 2021). This structure positions private stewardship as a key mediator of species occurrence and biodiversity patterns, in which fine-scale variation in management decisions can generate divergent ecological outcomes across the landscape.

In this study, we integrated historical and contemporary datasets to examine how long-term LULC change has reshaped vertebrate distribution ranges across Texas. Using Vernon Bailey’s late nineteenth-century surveys as a historical benchmark, we quantified changes in the distributions of 100 terrestrial and semi-aquatic vertebrate species over nearly two centuries of landscape transformation. To disentangle the drivers of these patterns, we asked how the magnitude and spatial patterns of LULC change have varied across Texas since the mid-nineteenth century, how these landscape changes were associated with patterns of species range expansion, contraction, and redistribution, and the extent to which habitat specialization and taxonomic identity predicted differential responses to landscape transformation, how patterns of range dynamics differed among major vertebrate groups, whether these differences reflect underlying variation in species traits, and how regional environmental gradients and ecoregional context interacted with land-use change to shape patterns of species distribution dynamics.

Consistent with these questions, we hypothesized that LULC change has been spatially heterogeneous, with agricultural expansion and urbanization driving the most pronounced transformations; that species traits mediate responses to landscape change, with habitat specialists exhibiting greater range contractions than generalist and highly dispersive species; that taxonomic groups exhibit predictable differences in range dynamics, with birds showing greater expansions, mammals intermediate responses, and reptiles and amphibians showing higher contraction rates; and that variation in species responses reflects interactions among land-use change, species traits, and regional environmental conditions. By linking long-term landscape transformation to species distribution dynamics, this study provides a century-scale perspective on how historical and ongoing habitat change have impacted vertebrate distribution ranges in heterogeneous landscapes. More broadly, it advances understanding of the mechanisms shaping biodiversity responses to global change and highlights the importance of historical baselines for predicting future ecological trajectories.

## 2. Methods

### 2.1 Land-use land-cover change

#### 2.1.1 LULC Datasets

We quantified LULC change in Texas from the mid-nineteenth century (1850) to the present (2023) using three primary datasets (Figure 1). We selected 1850 as the historical baseline because it precedes the first systematic biological survey of Texas (1889–1905) conducted by Vernon Bailey and represents a pre-survey landscape context (Bailey, 1905). This period (1850 onwards) also coincides with the onset of rapid anthropogenic land-cover change across the region (Afanador & Kjelland, 2025; Doughty, 1986). We used a broad temporal dataset spanning 1850-2020 at 1 km resolution, reconstructed from historical records, model-based LULC products, and census data by X. Li et al. (2023).

**Figure 1.**
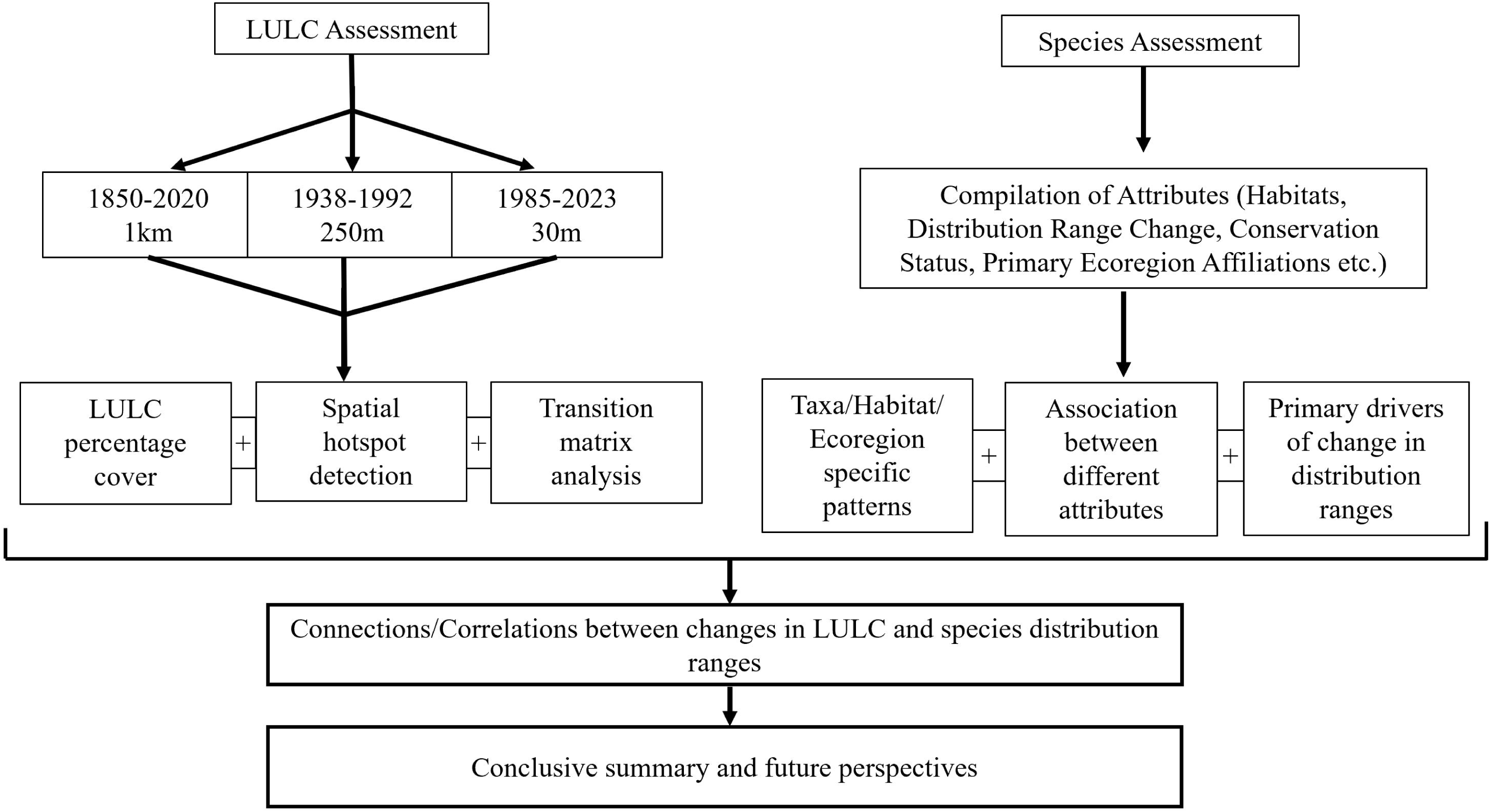
Overview of the methodological framework implemented in this study

Historical LULC reconstructions carry inherent uncertainties due to incomplete records, model assumptions, and differences in spatial resolution. To account for these uncertainties, we used multiple complementary datasets spanning different temporal resolutions and sources, harmonized classifications across datasets, and focused on broad-scale trends and transitions, thereby emphasizing patterns robust to potential errors in any single dataset. Accordingly, we incorporated two complementary finer-scale land-cover datasets: (1) a modeled historical dataset representing 1938–1992 at 250 m resolution (Sohl et al., 2018), and (2) an observed satellite-based dataset covering 1985–2023 at 30 m resolution (U.S. Geological Survey, 2024). Both were obtained from the Annual National Land-cover Database. The 1850–2020 dataset originally comprised 10 land-cover classes, whereas the other two datasets contained 16 classes (Details in Table S1). To ensure comparability across time and scale, we reclassified and merged some land-cover categories in the finer-resolution datasets to match the 10-class scheme used in 1850-2020 dataset. All analyses were therefore conducted using this unified 10-class classification for Texas. Although annual data were available for all three datasets, we selected data at 5-year interval to balance temporal resolution with computational efficiency.

#### 2.1.2 Temporal trends and transitions in LULC classes

For each dataset, we calculated the proportional area covered by each LULC class and assessed temporal trends in each class by comparing their area proportions across all time periods. To examine landscape transformation beyond net gains and losses, we constructed transition matrices that quantify the direction and magnitude of land-cover conversions between classes across successive intervals. To evaluate spatial variation, we summarized LULC composition and transitions across the 10 ecoregions of Texas between the initial and final timepoints of the three datasets. All the trend analysis and transition matrices computations were performed in R version 4.33. The final transition map of Texas was generated using Circos viewer (Krzywinski et al., 2009), a circular data visualization tool well-suited for representing transition matrices (Chen et al., 2017; Hu et al., 2020; Kosti et al., 2011).

#### 2.1.3 Spatial Hotspot Extraction

To identify statistically significant clusters of land-cover change, we calculated the Getis-Ord Gi* statistic (Getis & Ord, 1992). This metric computes a z-score for each pixel, reflecting the change in that location and among its neighbors relative to the global mean. On this scale, high positive z-scores indicate hotspots, while high negative z-scores identify cold spots. Statistical significance was assessed using associated p-values, with thresholds (e.g., p<0.01) to filter meaningful hotspots and cold spots. To reduce computational burden while retaining ecological relevance, all LULC datasets were resampled to a common 2.5 km² grid. This scale corresponds to mean spatial extents over which many vertebrate species respond to landscape change (Andrén, 1994; Provost et al., 2020; Solem et al., 2025; Williams et al., 2022). Hotspot analyses were performed in R by utilizing the local G function from the spatial R package called “spdep” (Bivand, 2025).

#### 2.1.4 Characterizing changes in Bailey’s survey locations in each ecoregion

We georeferenced 190 locations from Bailey’s Texas survey (1889–1905; Bailey, 1905) using archival maps reproduced by Schmidly et al (2022). Scanned maps were imported into ArcGIS Pro and aligned to a modern Texas base map using stable landmarks such as rivers, towns, and county boundaries. Sampling sites were then digitized as point features and assigned geographic coordinates. Where exact modern matches were unavailable, locations were approximated using the closest historical geographic references and verified against Bailey’s original locality descriptions. Getis-Ord Gi* values were extracted for each location to classify local land-cover change as hotspot, cold spot, or non-significant. Aggregating the proportion of points within each LULC change class by ecoregion allowed us to assess whether Bailey’s historical sampling locations experienced disproportionate land-cover change across regions.

### 2.2 Vertebrate distribution range changes

#### 2.2.1 Species selection and occurrence data

We assessed changes in the distribution ranges of 100 native vertebrate species (except fish) across Texas to evaluate long-term responses to LULC change (Figure 1; Table S2). Though we initially compiled a large pool of vertebrate species, we retained only those with robust and validated information, excluding taxa with uncertain records. Our aim was not to comprehensively represent the entire Texas vertebrate fauna, but to construct a representative dataset of each vertebrate class suitable for long-term, century-scale analyses. Vertebrates were selected because century-scale analyses require extensive historical baselines and relatively well-documented occurrence and trait data (Donaldson et al., 2016; Oliver et al., 2021; Schmitt et al., 2018; Titley et al., 2017). Biodiversity databases and studies are often biased towards vertebrates, making them a useful group for comparing spatiotemporal trends statewide (Donaldson et al., 2016; Titley et al., 2017). Species were selected to span four vertebrate classes (20 amphibians, 30 birds, 30 mammals, 20 reptiles), a range of body sizes, habitat affinities, and varying sensitivities to landscape modification including species reported to be positively or negatively affected by land use and land cover change. Collectively, they cover around 13% of all Texan vertebrates (except fish) and occupy most of Texas’ geographic extent, capturing high environmental and ecological heterogeneity across multiple ecoregions (Holt et al., 2020). We defined a species’ distribution range as the geographic area across which it occurs across Texas, encompassing all known populations within different ecoregions, counties and habitats, whereas changes in distribution range referred to expansions, contractions, or shifts in occupied geographic area over time.

Historical species occurrence data were primarily drawn from Vernon Bailey’s First Biological Survey of Texas (1889–1905), the earliest systematic documentation of the state’s mammalian and reptilian fauna and ecological conditions (Bailey, 1905; Schmidley et al., 2022). Historical species records carry uncertainties from sampling gaps and uneven coverage; we mitigated this by aggregating multiple sources and focusing on broad, long-term distributional trends across Texas. Additional records were obtained from Texas Nature Parks, Texas A&M University’s Biodiversity Research and Teaching Collection (BRTC), Vertebrate Database of Museum at Texas Tech University and the Global Biodiversity Information Facility (GBIF), covering all vertebrates except fish. For species not documented prior to 1890, distribution range change was assessed from their earliest available records to the present. For each species, we compiled the following attributes: scientific name, taxonomic group (Mammalia, Aves, Reptilia, Amphibia), primary habitat (mapped to LULC classes), distribution range change (increase, decrease, or range shift), primary ecoregions occupied (single, two, or multiple), and conservation status according to International Union for Conservation of Nature IUCN) and Texas Parks and Wildlife Department (TPWD) with IUCN categories collapsed into Threatened and Near Threatened/Least Concern (NTLC), and TPWD categories into state listed (Threatened/Endangered) and non-listed. Species were classified as having increasing distribution ranges if at least one county in Texas (without major further loss in any county) was added in the current distribution but absent from the historical range. Decreasing ranges were defined as those in which at least one county present in the historical distribution was absent from the current records (without major further addition in any county). Range shifts were defined as changes in species distribution between historical and contemporary records involving both gains and losses of counties, indicating spatial redistribution rather than unidirectional expansion or contraction. Species that had 70% of their distribution range consisting of only one habitat were considered habitat specialists, otherwise generalists (O’Reilly et al., 2022). Habitat associations among the 100 species were: barren (6), forest (30), grassland (15), shrubland (14), wetland (16), and generalist (18) (Details in Table S2).

#### 2.2.2 Species Distribution Range Change

To assess multivariate patterns influencing species distribution range changes, we examined how these changes (increase/decrease/shift) were related to taxonomic groups, primary habitat, and ecoregion association, and finally, how these relationships corresponded with observed land-cover changes in Texas. We first summarized species attributes and distributional patterns to provide an overview of the dataset. Associations among different attributes were quantified using Cramér’s V implemented in the R package vcd (Meyer et al., 2015).

To represent the latent structure among species attributes/variables in a reduced-dimensional space, we performed a Multiple Correspondence Analysis (MCA) using the FactoMineR package and visualized outputs with factoextra (Lê et al., 2008). MCA is appropriate for categorical datasets and is based on chi-square distances (Abdi & Valentin, 2007). To reduce sparsity, the Primary Ecoregion category was excluded, and distribution range change was limited to Increase and Decrease categories (as range shifts were rare).

Further, we evaluated the predictive power of species and habitat attributes on distribution range changes using a Random Forest (RF) classification model (R package *randomForest*) (Breiman, 2001). Random Forest is a non-parametric ensemble approach robust to multicollinearity, complex interactions, and mixed predictor types, including categorical variables (Cutler et al., 2007; Strobl et al., 2008). In addition to the species-level covariates described above, we incorporated ecoregion-specific LULC change metrics, including changes in percentage cover (Δ percentage cover) and landscape fragmentation (Δ fragmentation) between 1850 and 2020 for each of the 10 LULC classes. Landscape fragmentation was quantified using changes in Euclidean Nearest Neighbor (ENN) distance, calculated as the distance between the centroids of each patch and its nearest neighboring patch within the same LULC class. (Bosch, 2019). All Δ metrics were extracted from the single occupied ecoregion for restricted species, averaged across the two occupied ecoregions for species occurring in exactly two ecoregions, and averaged across all ecoregions for generalist species. Only for wetland and barren areas Δ changes were conducted between 1938 and 1992 due to limitations in the historical reconstruction of these land-cover types in the 1850-2020 dataset (X. Li et al., 2023). Ecoregions represented by fewer than two species were pooled to ensure model stability. Models were trained using 500–10,000 trees, with the optimal number determined by out-of-bag (OOB) error rates. Predictor importance was quantified using Mean Decrease in Accuracy (MDA), measuring the reduction in classification accuracy following variable permutation, and Mean Decrease in Gini (MDG), reflecting each predictor’s contribution to node purity across the forest.

## 3. Results

### 3.1 Statewide LULC trends and species distribution range changes

Over nearly two centuries, Texas transitioned from landscapes dominated by native and semi-natural vegetation (forest, grassland, shrubland, and wetland combined; hereafter natural cover) to systems increasingly structured by human land-use, with distinct phases captured by the different LULC datasets (Figures 2, S1, and S2). At the broadest scale, the 1850–2020 period is characterized by a decrease in natural cover from ∼ 97% to 71% statewide, while cropland, pasture, and urban land expanded to 26% (Figures 2 and S3). Statewide transition matrices further revealed substantial conversions of natural habitats, including 23% of forests converted to pasture and 24% of grasslands converted to cropland over the last 170 years. Although cropland expansion peaked in the late 1800s, finer-resolution datasets (1938–1992 at 250 m and 1985–2023 at 30 m) revealed a subsequent decline of ∼10% in croplands between 1938 and 2023, accompanied by continued declines in natural cover (Figures S3, S4, and S5). Over the past ∼80 years, land-cover structure has been characterized by widespread reconfiguration, with ∼65% of cropland converted to shrubland, forest, pasture, or urban land. Notably, shrubland encroachment increased, particularly into grasslands (13%) and former croplands (10%) (Figures S8, S9, and S10).These long-term LULC changes correspond with marked shifts in the distribution of mammals, birds, reptiles, and amphibians. Across the 100 vertebrate species assessed, range contractions were the dominant response, occurring in 68% of species (e.g., Timber Rattlesnake, Pronghorn), whereas 28% showed range expansions (e.g., Blue Jay, Texas Toad), and only 4% exhibited range shifts (American Badger, Common Muskrat, Mule Deer, and Northern River Otter). Despite widespread distributional changes, only 8% of species are classified as Threatened by the IUCN, and 21% are classified as Threatened/Endangered by the Texas Parks and Wildlife Department (TPWD).

**Figure 2.**
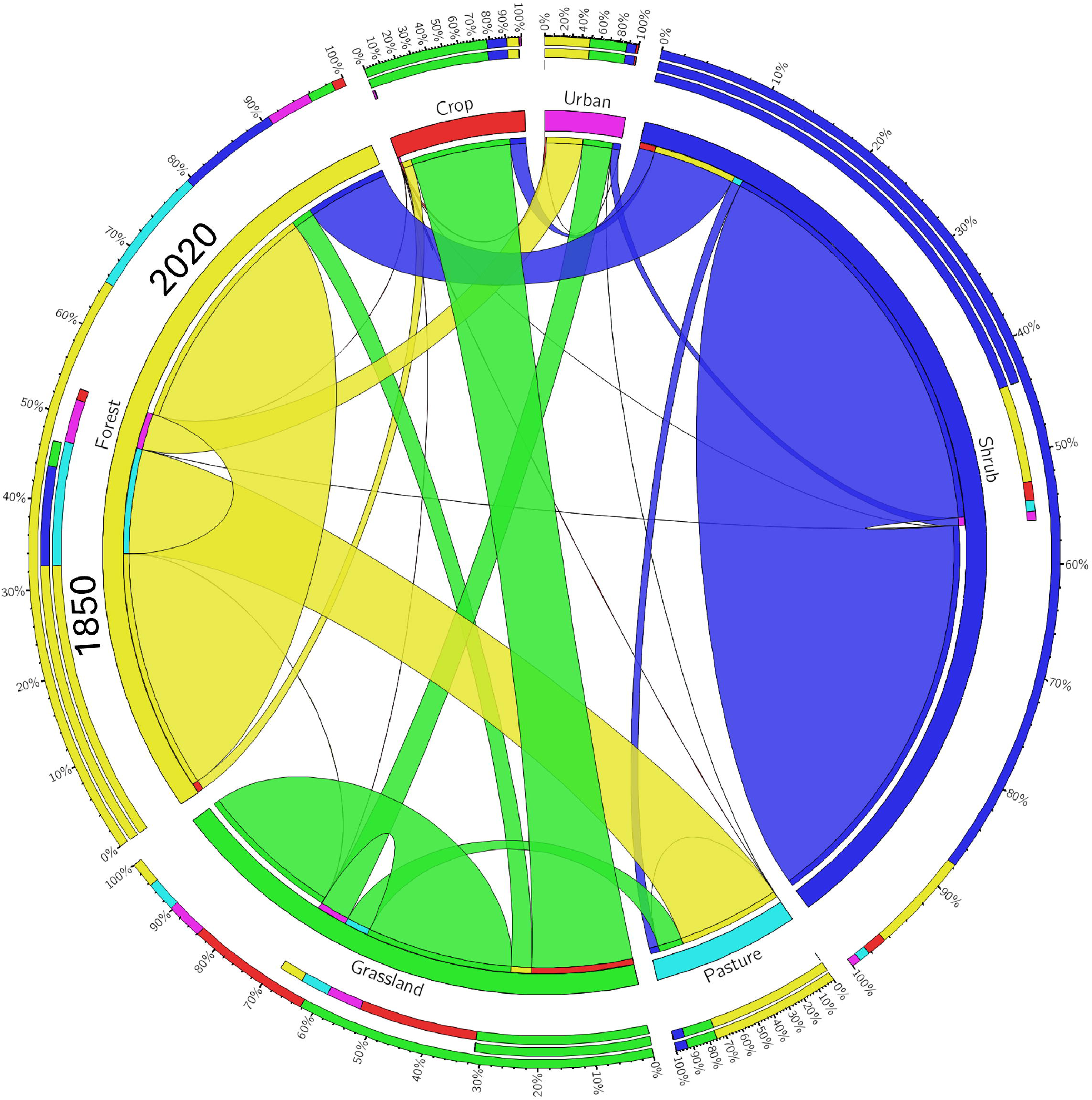
Chord diagram illustrating land-use and land-cover (LULC) transitions between 1850 and 2020. Directionality of change is indicated by ribbons flowing from their closed ends (1850) to their open ends (2020). Three concentric outer rings summarize the magnitude of transitions for each LULC class: the innermost ring shows the total area transitioning *from* a class, the middle ring shows the total area transitioning *to* a class, and the outer ring shows the total area involved in all transitions associated with that class.

Clear taxonomic differences emerged. Range contractions were most pronounced among amphibians (80%; N = 16/20; e.g., Great Plains Toad) and reptiles (95%; N = 19/20; e.g., Ornate Box Turtle). In contrast, range expansions were more common among mammals (40%; N = 12/30) and birds (36.7%; N = 11/30). These patterns suggest potential differences among vertebrate groups in their responses to long-term landscape change, associated with variation in dispersal capacity, physiological tolerance, and sensitivity to habitat fragmentation.

Habitat specialization further structured species responses (Table 2). Species associated with declining or heavily modified habitats experienced substantially higher contraction rates than generalists, particularly wetland (81%; N = 13/16; e.g., Swamp Rabbit), shrubland (78%; N = 11/14; e.g., Texas Horned Lizard), grassland (73%; N = 11/15; e.g., Texas Kangaroo Rat), and forest (68%; N = 20/30; e.g., Red-cockaded Woodpecker) taxa. In contrast, generalist species showed lower contraction rates (36%; N = 7/19; e.g., Ornate Tree Lizard). These results highlight habitat specialization as a key axis along which species responses to land-use change diverge. Trait-by-habitat interactions revealed additional nuance to these trends. Among habitat specialists, herpetofauna were disproportionately affected, with 92% of specialist amphibians and reptiles contracting (N = 33/36), including complete contraction of wetland-associated species (100%; N = 7/7; e.g., Houston Toad). Within birds, grassland specialists (100%; N = 3/3; e.g., Lesser Prairie-Chicken) and wetland specialists (100%; N = 4/4; e.g., Reddish Egret) exhibited the highest range contractions, whereas mammals showed the strongest declines in shrubland habitats (100%; N = 3/3; e.g., Ocelot). In contrast, localized range expansions were observed among forest-associated mammals (80%; N = 4/5; e.g., Virginia Opossum) and shrubland-associated birds (100%; N = 2/2; e.g., Common Ground Dove). Taken together, these observations underscore that species responses to long-term landscape change are shaped by the taxonomic identity, habitat specialization, and trait-mediated ecological flexibility (e.g., dispersal ability and habitat breadth).

### 3.2 Spatial heterogeneity in LULC change and species responses across ecoregions

Getis-Ord Gi* hotspot analysis over the 170-year period showed that the most pronounced LULC changes occurred across the areas between cities of Dallas, Houston, and San-Antonio (Table 1, Figure 3 left panel). Among ecoregions, the Post Oak Savannah (76% of all pixels) and Blackland Prairies exhibited the highest proportion of hotspot areas (64% of all pixels), whereas Trans-Pecos showed minimal change (less than 1% of all pixels; Figures 3A and S6). In recent decades, many areas within the Post Oak Savannah and Blackland Prairies exhibited predominantly non-significant change (89% and 91% of pixels; Figures 3B and 3C), suggesting that these regions have reached a relative state of stabilization. In contrast, the Piney Woods emerged as a contemporary hotspot of change, with 67% of the pixels classified as hotspots between 1985-2023 (Figures 3C, S11 - S13). Analysis of Bailey’s historical survey locations revealed a similar spatial pattern, with the highest proportion of significant hotspots of change occurring in the Blackland Prairies, Piney Woods, and Post Oak Savannah (Figure S7).

**Figure 3.**
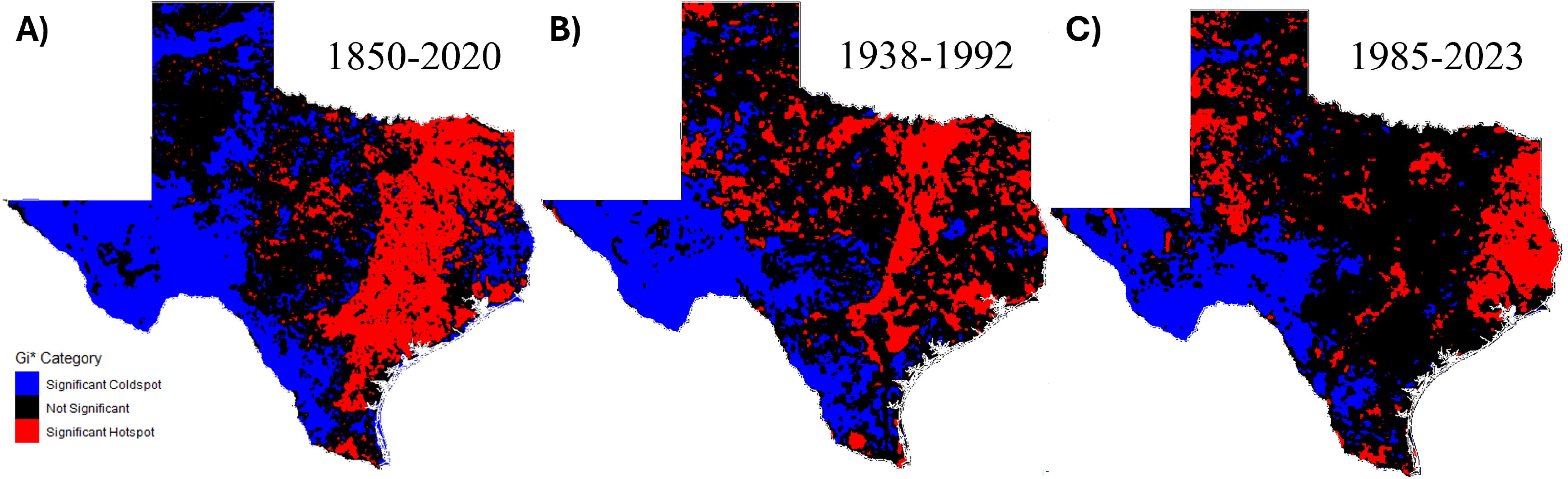
Spatial hotspots and coldspots of LULC change across the three temporal datastets. A) 1850-2020 B) 1938-1992, and C) 1985-2025.

**Table 1.** Summary of dominant land-cover classes and their changes in Texas ecoregions across three temporal datasets. For each ecoregion, the table reports the starting and ending dominant class, percent cover, major land-cover transitions, and the percentages of pixels identified as significant hotspots and cold spots of change

| Ecoregion Name | Dominant Class<br>Name_Start<br>year | % cover of<br>Dominant Class<br>class_Start year | Dominant Class<br>Name_End year | % cover of<br>Dominant<br>class_End year | Dominant<br>Transitions | Significant<br>Change Hotspot<br>pixel % | Significant<br>Change Coldspot<br>pixel % |
| --- | --- | --- | --- | --- | --- | --- | --- |
| 1850-2020(1km) |  |  |  |  |  |  |  |
| Blackland Prairie | Forest | 50 | Pasture | 33 | Forest-Pasture | 64.4 | 0.83 |
| Cross Timbers | Forest | 61 | Forest | 69 | Crop-Forest | 18.14 | 11.26 |
| Edwards Plateau | Shrub | 77 | Shrub | 53 | Shrub-Forest | 3.51 | 43.72 |
| Gulf Prairies | Grassland | 55 | Pasture | 22 | Grassland-Pasture | 41.77 | 13.78 |
| High Plains | Grassland | 67 | Crop | 43 | Grassland-Crop | 5.09 | 20.12 |
| Piney Woods | Forest | 81 | Forest | 55 | Forest-Pasture | 29.57 | 24.75 |
| Post Oak Savanah | Forest | 85 | Pasture | 58 | Forest-Pasture | 75.86 | 1.03 |
| Rolling Plains | Shrub | 53 | Shrub | 44 | Grassland-Crop | 5.51 | 35.68 |
| South Texas Plains | Shrub | 62 | Shrub | 45 | Shrub-Forest | 17 | 37.44 |
| Trans-Pecos | Shrub | 87 | Shrub | 83 | Shrub-Forest | 0.1 | 86.92 |
| 1938-1992(250m) |  |  |  |  |  |  |  |
| Blackland Prairie | Crop | 35 | Pasture | 42 | Crop-Pasture | 64.22 | 0.03 |
| Cross Timbers | Grassland | 28 | Grassland | 32 | Crop-Grassland | 27.05 | 2.83 |
| Edwards Plateau | Shrub | 53 | Shrub | 56 | Crop-Shrub | 5.5 | 45.09 |
| Gulf Prairies | Crop | 23 | Pasture | 21 | Pasture-Crop | 23.36 | 5.79 |
| High Plains | Crop | 57 | Crop | 52 | Crop-Grassland | 15.03 | 17.23 |
| Piney Woods | Forest | 53 | Forest | 66 | Crop-Forest | 23.74 | 4.76 |
| Post Oak Savanah | Pasture | 38 | Pasture | 39 | Crop-Forest | 25.92 | 4.26 |
| Rolling Plains | Grassland | 36 | Grassland | 41 | Crop-Shrub | 16.03 | 4.81 |
| South Texas Plains | Shrub | 53 | Shrub | 54 | Crop-Shrub | 9.59 | 36.57 |
| Trans-Pecos | Shrub | 73 | Shrub | 72 | Shrub-Crop | 0.52 | 87.51 |
| 1985-2023(30m) |  |  |  |  |  |  |  |
| Blackland Prairie | Pasture | 35 | Pasture | 36 | Crop-Pasture | 9.39 | 2.07 |
| Cross Timbers | Grassland | 43 | Grassland | 40 | Crop-Grassland | 8.57 | 0.41 |
| Edwards Plateau | Shrub | 83 | Shrub | 85 | Shrub-Forest | 3.42 | 47.71 |
| Gulf Prairies | Pasture | 28 | Pasture | 28 | Crop-Pasture | 12.35 | 2.67 |
| High Plains | Crop | 57 | Crop | 46 | Crop-Grassland | 28.03 | 2.84 |
| Piney Woods | Forest | 47 | Forest | 42 | Shrub-Forest | 79.08 | 0.07 |
| Post Oak Savanah | Pasture | 50 | Pasture | 51 | Crop-Pasture | 7.5 | 0.59 |
| Rolling Plains | Shrub | 52 | Shrub | 61 | Crop-Shrub | 5.5 | 7.95 |
| South Texas Plains | Shrub | 63 | Shrub | 65 | Pasture-Shrub | 6.18 | 21.62 |
| Trans-Pecos | Shrub | 91 | Shrub | 92 | Grassland-Shrub | 2.15 | 67.19 |

**Table 2.** Summary of species by taxonomic group and primary habitat, showing the number of species, the percentage of species within each taxonomic group occurring in a particular habitat, the proportion of species in each taxon–habitat combination with decreasing or increasing distributions, observed range shifts, and conservation status according to IUCN and TPWD listings

|  | **Primay**<br>**Habitat** | **Number**<br>**of**<br>**species** | **% of**<br>**taxa**<br>**type**<br>**in**<br>**habitat** | **%**<br>**decreasing** | **%**<br>**increasing** | **%range_shift** | **%**<br>**IUCN_threatened** | **%**<br>**TPWD_listed** |
| --- | --- | --- | --- | --- | --- | --- | --- | --- |
| Amphibia | Forest | 9 | 45 | 77.78 | 22.22 | 0 | 11.11 | 22.22 |
|  | Generalist | 2 | 10 | 50 | 50 | 0 | 0 | 50 |
|  | Grass | 2 | 10 | 50 | 50 | 0 | 0 | 0 |
|  | Shrub | 2 | 10 | 100 | 0 | 0 | 0 | 0 |
|  | Wetland | 5 | 25 | 100 | 0 | 0 | 20 | 20 |
| Aves | Barren | 2 | 6.67 | 100 | 0 | 0 | 0 | 0 |
|  | Forest | 12 | 40 | 66.67 | 33.33 | 0 | 8.33 | 33.33 |
|  | Generalist | 7 | 23.33 | 28.57 | 71.43 | 0 | 0 | 0 |
|  | Grass | 3 | 10 | 100 | 0 | 0 | 33.33 | 33.33 |
|  | Shrub | 2 | 6.67 | 0 | 100 | 0 | 0 | 0 |
|  | Wetland | 4 | 13.33 | 100 | 0 | 0 | 0 | 25 |
| Mammalia | Barren | 2 | 6.67 | 100 | 0 | 0 | 50 | 100 |
|  | Forest | 5 | 16.67 | 20 | 80 | 0 | 0 | 0 |
|  | Generalist | 8 | 26.67 | 25 | 62.5 | 12.5 | 0 | 12.5 |
|  | Grass | 7 | 23.33 | 57.14 | 28.57 | 14.28 | 14.29 | 14.29 |
|  | Shrub | 3 | 10 | 100 | 0 | 0 | 0 | 33.33 |
|  | Wetland | 5 | 16.67 | 40 | 20 | 40 | 0 | 0 |
| Reptilia | Barren | 2 | 10 | 100 | 0 | 0 | 50 | 50 |
|  | Forest | 4 | 20 | 100 | 0 | 0 | 25 | 50 |
|  | Generalist | 2 | 10 | 100 | 0 | 0 | 0 | 0 |
|  | Grass | 3 | 15 | 100 | 0 | 0 | 0 | 33.33 |
|  | Shrub | 7 | 35 | 85.71 | 14.29 | 0 | 14.29 | 42.86 |
|  | Wetland | 2 | 10 | 100 | 0 | 0 | 0 | 0 |

Species assessment at the ecoregion level reflected similar spatial patterns. Species inhabiting multiple ecoregions showed balanced outcomes, with equal proportions of range decreases and increases (N=48, 50%). In contrast, 91% of species primarily found in the Piney Woods declined (N=11/12; e.g., Red Cockaded Woodpecker). Similarly, 90% of species in the Trans-Pecos showed range contractions (N=8/9; e.g., Dunes Sagebrush Lizard), despite relatively limited LULC change in this region. The South Texas Plains ecoregion had the highest number of TPWD-state-listed endangered or threatened species (N=5/8, 62.5%; e.g., ocelot).

### 3.3. Drivers and multivariate structure of species distribution change

Pairwise associations among covariates revealed several strong relationships (Table S3). The highest associations were observed between IUCN and TPWD conservation status (V = 0.55, p < 0.001), IUCN status and primary ecoregion (V = 0.58, p < 0.001), and primary ecoregion and primary habitat (V = 0.56, p < 0.001). Taxonomic group showed moderate but significant association with both distribution range shift (V = 0.32, p < 0.001), and IUCN status (V = 0.35, p = 0.02). Other variable pairs had lower Cramér’s V values and were not statistically significant (Table S3).

Multivariate analysis further revealed gradients structuring species responses (Figure 4). The first two dimensions of the MCA explained 18.2% and 12.2% of total inertia/variance, respectively, capturing the dominant gradients in the data. In the ordination space, species exhibiting range increases tended to cluster near generalists, whereas reptiles and amphibians were more often associated with range contractions. Species listed as Least Concern or Near Threatened by IUCN were generally not listed under TPWD, reflecting concordance between global and state-level assessments. Among habitat types, grasslands, shrublands, forests, and wetland species were more often associated with range reduction.

**Figure 4.**
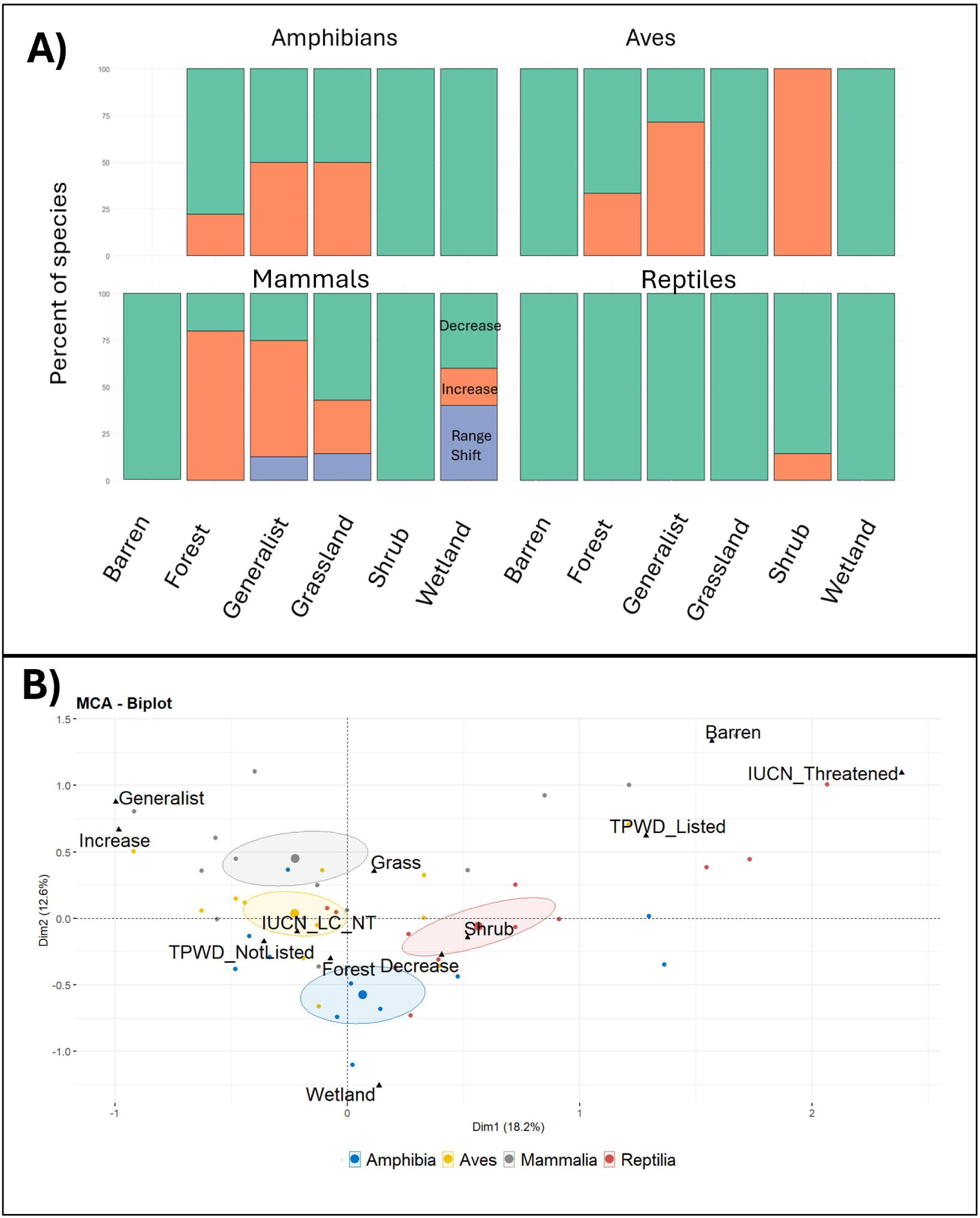
(A) Stacked bar plots showing the proportion of species exhibiting range contraction, expansion, or spatial shifts within each primary habitat category for amphibians, birds, mammals, and reptiles. (B) Multiple correspondence analysis (MCA) biplot summarizing associations among range-change categories (increase/decrease), primary habitats, and conservation status. Points are colored by taxonomic group with ellipses highlighting group-level tendencies in multivariate space.

RF classification provided a complementary assessment of species attributes and landscape variables. Model performance stabilized at ∼5,000 trees based on OOB error saturation, and this value was used in the final model. The RF model yielded an OOB rate of 32.32%, indicating moderate discriminatory power in classifying species into increasing, decreasing, and range-shift categories. Variable importance metrics (Mean Decrease in Accuracy and Mean Decrease in Gini) identified primary habitat, vertebrate class, and ecoregion as the strongest predictors of distribution range change (Figures 5A and 5B). Changes in urban area cover and wetland fragmentation also contributed meaningfully to model performance, although to a lesser extent (Figure 5A). In contrast, conservation status (IUCN and TPWD) had minimal influence on RF model predictions (Figure 5A).

**Figure 5.**
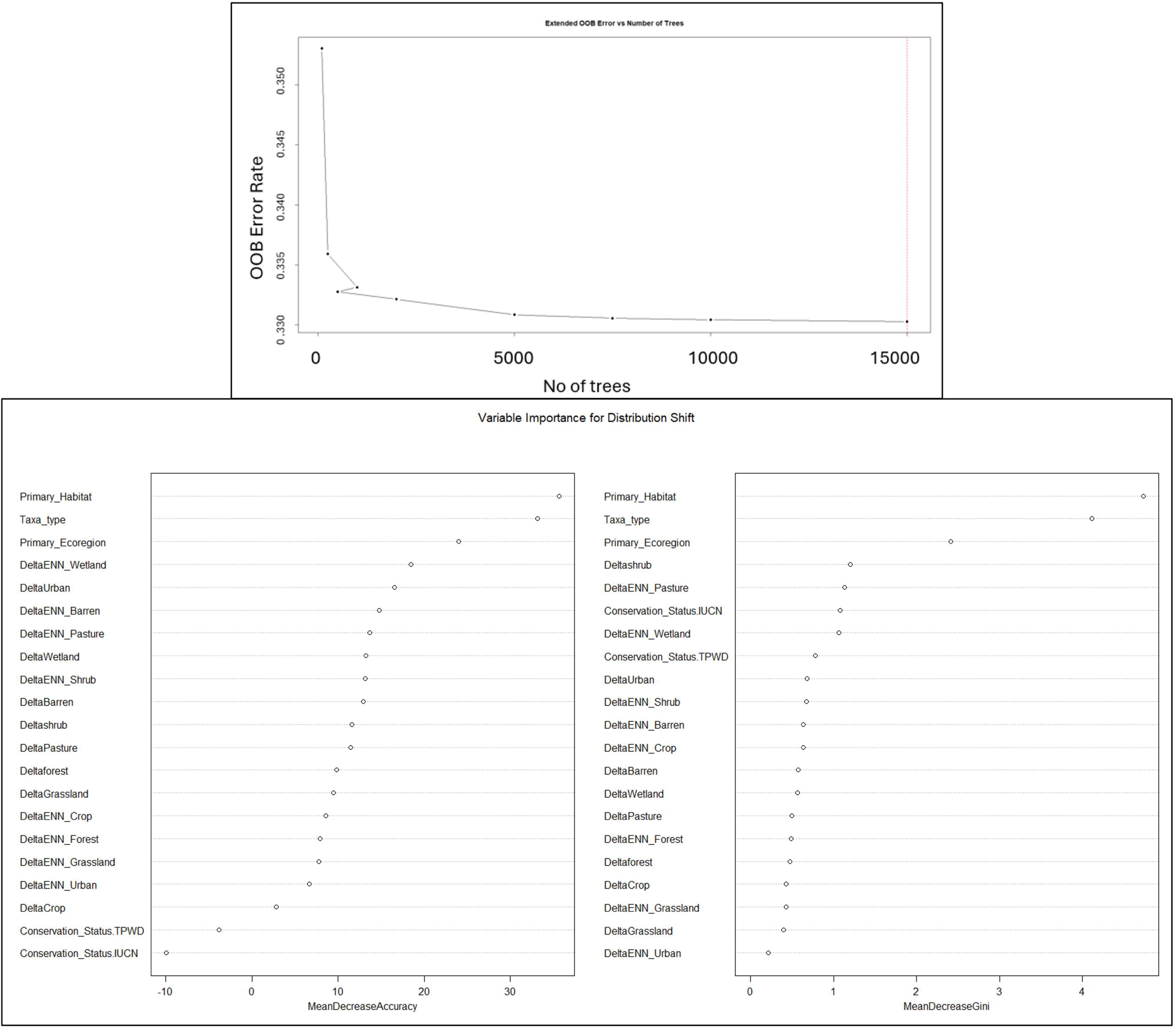
Random forest model performance and variable importance for explaining species distribution range changes. (A) shows the out-of-bag (OOB) error rate as a function of the number of trees, indicating progressive error reduction and stabilization as model complexity increases. (B) displays variable importance ranked by mean decrease in accuracy (left) and mean decrease in Gini index (right), with predictors ordered from the strongest to weakest contributors in explaining distribution range change across species.

## 4. Discussion

Our results show that nearly two centuries of LULC change in Texas have contributed to pronounced and spatially uneven restructuring of vertebrate species distributions. While long-term LULC change sets the landscape template, species responses are primarily governed by ecological filtering through habitat affinity, taxonomic traits, and ecoregion. Viewed through a community-assembly lens, these patterns are consistent with strong ecological filtering whereby landscape transformation favored generalist and mobile taxa while constraining habitat specialists with narrower ecological requirements. Across the state, these dynamics are amplified in regions functioning as biogeographic crossroads where strong environmental gradients, overlapping ecoregions, and high species turnover allow divergent ecological responses to emerge within a single landscape (Lombardi et al., 2020; Schmidley et al., 2022). As a result, long-term LULC change across Texas has not produced uniform biodiversity loss, but rather a mosaic of range expansions, contractions, and redistributions shaped by the combined effects of land-use history, environmental context, and species-specific ecological traits. These patterns are consistent with broader evidence that biodiversity responses to environmental change are spatially heterogeneous and shaped by species traits and environmental context, not solely by the extent of LULC changes, with responses varying across environmental gradients.

### 4.1 Long-term LULC trajectories and regional context

Early land conversion in Texas was dominated by cropland expansion and pasture development during the late nineteenth and early twentieth centuries. In contrast, more recent trajectories are characterized by declining cropland extent, shrub encroachment, secondary forest regeneration, and rapid urban growth- patterns independently documented across Texas landscapes (Asner et al., 2003; Cain & Shelton, 2001; X. Li et al., 2023; Olariu et al., 2024; Ritchie, 2025; A. Smith et al., 2025; Wilcox & Huang, 2010; Wilkins et al., 2009). These shifts reflect distinct phases of ecological filtering, in which changes in land-use practices altered the availability and configuration of habitats, differentially influencing species persistence across taxa both within and between regions.

The statewide trajectories are disproportionately associated with a subset of ecoregions that differ in their intensity of land-use changes. Among all ecoregions, the Blackland Prairies and Post Oak Savannah experienced the earliest and most intensive changes between 1850 and 2020, confirmed through hotspot analyses. Most of the Blackland Prairie prairies were cleared for cotton cultivation (46% loss in grasslands between 1850-2020), making it one of the most endangered ecoregions in North America (Diamond & Smiens, 1993; Griffith et al., 2007; Schmidley et al., 2022). In the Post Oak Savannah, native grasslands declined by 80% over the same period, as widespread clearing and cultivation produced landscapes dominated by dense pasture grasses, woody undergrowth, and thicketized woodlands (Griffith et al., 2007; Olariu et al., 2024; Rideout, 1994). In contrast, the Trans-Pecos ecoregion has remained relatively intact, retaining much of its native desert and shrubland owing to arid conditions, low population density, and limited agricultural and urban pressure (Griffith et al., 2007; TPWD, 2020). More recently, the Piney Woods have emerged as a new hotspot of change, associated with forest fragmentation, urban development, poor regeneration, fire suppression, and declines in evergreen and mixed forests (6% and 25% loss, respectively, between 1985 and 2023) (Comer & Steumke, 2011; Texas A&M Forest Service, 2009; Wall et al., 2019). Together, these contrasts highlight how baseline environmental context shapes the influence of land-use change across ecoregions and reinforce the expectation that LULC change operates through region-specific pathways.

### 4.2 Trait-mediated ecological filtering of species distributions

The dynamic LULC changes observed across Texas were also accompanied by variable responses in the 100 vertebrates observed in this study. Importantly, species selection was based on data availability and historical coverage across taxa and habitat types, rather than observed range change outcomes, ensuring that the observed patterns are not an artefact of sampling design.As expected, species tied to shrinking or heavily modified habitats exhibited the most pronounced range contractions, with habitat specialists (wetlands, grasslands, shrublands, and forests) declining more than generalist species, consistent with their heightened vulnerability to habitat loss and fragmentation (Clavel et al., 2011). Wetland specialists showed the strongest contractions (81% of species), consistent with the well documented susceptibility of wetlands globally to drainage, hydrological alteration, pollution, and rapid loss due to agricultural and urban expansion (Robertson et al., 2025). Grassland, shrubland, and forest specialists also declined in line with pervasive habitat conversion and fragmentation across these ecosystems (Clavel et al., 2011).

The magnitude and direction of distribution range changes differed among vertebrate groups, largely consistent with expectations. Mammals and birds exhibited similar proportions of range increase (40% and 37% of species, respectively), whereas reptiles and amphibians showed the lowest levels of range expansions (5% and 20% of species, respectively), reflecting their limited dispersal ability, sensitivity to habitat fragmentation, narrow thermal tolerances, and dependence on microhabitat conditions (Colino-Rabanal & Lizana, 2012; Dodd, 2010; Todd & Rothermel, 2006; Veselka et al., 2023). These patterns are broadly consistent with observations from other heterogeneous landscapes (Beja & Alcazar, 2003; Biancolini et al., 2024; Newbold et al., 2019; Pacifici et al., 2017; Subba et al., 2018) and global assessments of vertebrate decline, particularly herpetofauna (Cox et al., 2022; Hayes et al., 2010; Q. Li et al., 2024; Luedtke et al., 2023; Subba et al., 2018). Within habitat specialists, wetland associated herpetofauna and birds showed major declines, while grassland species, particularly birds, exhibited shrinking distribution ranges consistent with broader continental trends (Correll et al., 2019; Douglas et al., 2023; Fern et al., 2023; Mallet et al., 2023; North American Bird Conservation Initiative, 2025; Rosenberg et al., 2019). Divergent responses among taxa occupying similar habitats further emphasize the role of species traits. For example, shrubland-associated mammal species exhibited range contractions, whereas some shrubland-associated bird species expanded their ranges, likely reflecting differences in mobility and foraging ecology, which mediate how organisms track changing habitat structure (Burnett & Koprowski, 2024; Ceballos et al., 2010; Torre & Palau, 2023). Apparent increases in forest associated mammals may reflect interacting effects of variable land-cover dynamics (e.g., afforestation or woody encroachment), altered disturbance regimes (e.g., hunting practices or prescribed fires) and/or conservation actions.

Patterns of distribution range change also varied among ecoregions. Species associated with the Piney Woods declined in parallel with substantial forest loss over the past four decades (TPWD, 2006). Because only a few selected species were restricted exclusively to the Post Oak Savannah and none to the Blackland Prairies, ecoregion-specific inferences for these systems were limited. Interestingly, species associated with the Trans-Pecos ecoregion exhibited range contractions, despite minimal LULC change, suggesting that additional drivers, such as climate change, may independently contribute to species vulnerability (Yancey et al., 2023). Given that numerous species occupy multiple ecoregions and most exhibit generalist habitat preferences, caution is warranted when drawing conclusions from ecoregion-level analyses.

Association analysis and multivariate analysis reinforced these patterns. The moderate association between IUCN and TPWD conservation status indicates that global and regional threat assessments are largely consistent. Strong associations with primary ecoregions and habitats indicate that higher-risk species are concentrated in specific regions and habitats.

Taxonomic group category was moderately associated with both range change and conservation status, reflecting that amphibians and reptiles have experienced greater contractions and are more likely to be threatened at present than many bird and mammal species. Similarly, the MCA revealed underlying ecological gradients in which generalist species aligned with expanding distribution ranges, whereas reptiles, amphibians, and habitat specialists aligned with contracting distribution ranges. Consistent with these results, RF models identified primary habitat, vertebrate class, and ecoregion as the strongest predictors of distributional range change, emphasizing that ecological context and not just the magnitude of land-cover change governs species’ response. These variables likely capture underlying factors such as microclimate, disturbance history, connectivity, and land-use legacies that shape species responses over time (Esser et al., 2025; Mi et al., 2017; Simon et al., 2023). Although metrics of ecoregion change and fragmentation capture net land-cover shifts and isolation of habitat patches, they do not fully represent the habitat quality and timing, sequencing, or cumulative effects of other disturbances that shape long-term ecological outcomes (Foster et al., 2003; Turner, 2001). Urban expansion and wetland fragmentation nevertheless contributed meaningfully to model performance, reflecting recent anthropogenic pressures. Taken together, our study shows that baseline ecological context embedded in habitat and ecoregion identity exerts a stronger influence on long-term species persistence than net land-cover change alone, emphasizing the importance of characterizing regional context and habitat specificity when assessing vulnerability or planning conservation strategies (Bevan et al., 2025; Lindenmayer et al., 2023; Newbold et al., 2019; Pacifici et al., 2017).

### 4.3 Strengths and limitations

This study is among the few to integrate approximately 170 years of LULC reconstructions with multi-taxa vertebrate distribution ranges across an entire state spanning multiple ecoregions (X. Li et al., 2023). Historical datasets, such as Bailey’s early twentieth-century surveys, provide a critical baseline for understanding long-term trends in species distributions and habitat change and identifying vulnerable regions (e.g., Blackland Prairie and Post Oak Savannah). By combining these records with temporal LULC reconstructions, species traits, multivariate approaches, we provide a mechanistic understanding of how ecological context, habitat specialization, and taxonomic identity jointly shape species range contractions and expansions. Unlike studies focused on single habitats or short time frames, this work captures spatiotemporal heterogeneity in both landscape change and biodiversity responses, revealing emergent patterns, including shifting hotspots of land-use change and the disproportionate vulnerability of specific taxa.

Several caveats remain. Historical species distributions and LULC reconstructions carry uncertainties, particularly for earlier time periods. The study examined distribution range changes of 100 vertebrate species, which may not fully capture variation across complete communities or allow formal analyses of community reassembly. This limitation is further exacerbated using categorical distribution data rather than quantitative abundance measures, limiting the ability to capture subtle or fine-scale changes. Although our analyses identify strong associations between land-use change and species distribution dynamics, their correlative nature limits direct inference of causality, and multiple interacting drivers, including climate change and biotic interactions, likely contribute to observed patterns. The RF model showed moderate predictive accuracy, indicating that unmeasured variables may influence range changes.

Furthermore, invertebrates, plants, and aquatic taxa were outside the scope of this study, limiting broader generalization. Despite these limitations, the framework provides a long-term, multi-scale perspective linking landscape transformations to biodiversity outcomes.

### 4.4 Implications for conservation and transferable model frameworks

Building on this long-term baseline, future work should integrate historical and contemporary species distribution data to better assess how past and ongoing landscape transformations influence species’ capacity to respond to future change. This includes assessing community-level patterns. adaptive potential, encompassing genetic diversity and evolutionary capacity, which may determine population persistence under novel environmental pressures (Aitken et al., 2024; Vajana et al., 2022; Westaway et al., 2025).

Texas has experienced substantial landscape change alongside rapid population growth, with continued expansion of urban land-cover projected over the coming decades (Guo & Zhang, 2021; A. Smith et al., 2025). Understanding how these long-term projected changes will affect species distributions and their range limits requires integrating contemporary biodiversity data with historical records. Museum specimens, such as those housed at Texas A&M University and other regional collections, can provide critical quantitative information for reconstructing past distributions, evaluating genetic variation, and improving predictions of species’ responses to land-use and climate change (Meineke et al., 2018; Schmitt et al., 2018; Suarez & Tsutsui, 2004). Revisiting historical survey points, including those documented by Bailey, can reveal areas of past and ongoing shifts in community composition (Beller et al., 2020; Labay et al., 2011). These shifts can be used to formulate conservation plans for historically vulnerable landscapes (Beller et al., 2020; Duren et al., 2012).

By integrating long-term land-use trajectories, species traits, and regional environmental gradients within a single administrative unit, Texas further highlights the need to account for historical land-use legacies when designing evidence-based conservation strategies at the administrative unit level. For example, private land conservation programs across Texas have converted millions of acres from agricultural use to wildlife management while retaining tax incentives; these programs require ongoing evaluation as species’ geographic limits continue to shift under ongoing changes (Benavidez et al., 2021). Additionally, because Texas encompasses multiple ecoregional crossroads and functions as a single administrative unit, these findings provide a strong foundation for state-level conservation planning and policy development involving both government agencies and private landowners. Finally, species distributions do not always align neatly with ecoregion boundaries, emphasizing the importance of state-level approaches to effectively address both widespread and patchily distributed species (McDonald et al., 2005; J. R. Smith et al., 2018).

Landscapes composed of multiple biogeographic crossroads, like Texas, emphasize the need for systematic, multi-scale assessments of land-use and climate change, species distributions, and ecological and evolutionary variability. Globally, such heterogeneous systems (e.g., Cerrado–Caatinga–Chaco interface, Mediterranean Basin, Southeastern United States, Himalaya) are particularly valuable because they combine large spatial coverage with diverse land-use trajectories within a single region, enabling detection of divergent ecological responses including range contraction, expansion, and potential shifts in species boundaries that may be obscured in more homogeneous regions (Arslan et al., 2025; Beirão et al., 2017; Cartwright & Wolfe, 2016; Ferreira & Beja, 2013; Hanberry, 2020; Martins et al., 2022; Pereira et al., 2023; Quintas-Soriano et al., 2022; Thorne et al., 2022; Wambulwa et al., 2021; Werneck, 2011). Because responses can differ markedly among taxa, habitats, and regions, integrating species traits within environmental context, and long-term land-use history is essential for understanding species distributions and possible community reassembly. In this study, these patterns collectively demonstrate that LULC change is spatially heterogeneous, that species traits strongly mediate distributional responses, and that these responses are shaped by regional environmental context. By integrating these dimensions, this framework establishes a transferable model for quantifying how species distribution range changes, traits, and specialization respond to LULC change, and offers a foundation for proactive, evidence based conservation planning in other heterogeneous landscapes experiencing rapid change.

## Author Contributions

**Shrutarshi Paul**: conceptualization, data curation, formal analysis, investigation, methodology, project administration, software, validation, visualization, supervision, writing-original draft, writing-review and editing. **Vivianna Borzym**: data curation, investigation, methodology, visualization, writing-review and editing. **Heather Prestridge**: data curation, investigation, methodology, validation, supervision, writing-review and editing. **Wenzhe Jiao**: investigation, methodology, software, validation, visualization, supervision, writing-review and editing. **Mary Katherine Gonder**: conceptualization, data curation, investigation, methodology, project administration, validation, visualization, writing-original draft, writing-review and editing, funding acquisition, resources, supervision.

## Acknowledgement

We thank Texas A&M University for financial support. We are grateful to the Texas A&M Biodiversity Research and Teaching Collections Curators for making historical and contemporary ecological data publicly available. We thank Fabrice Kentatchime, Alvine Dadjo, and Dr. Kevin Njabo for assistance with data processing and constructing the methodological framework. We also acknowledge the technical assistance from Jiawei Wei in acquiring LULC data. Finally, we appreciate the constructive comments from reviewers and colleagues that greatly improved the clarity and scope of this work.

## Data Availability Statement

Additional data will be made available upon request or publication.

## Artificial Intelligence Statement

During the preparation of this manuscript, the authors used OpenAI ChatGPT to assist with language editing, manuscript organization, and development of a preliminary graphical abstract concept. All data collection, statistical analyses, interpretation, and final figure preparation were conducted by the authors without the use of AI-generated analyses. The authors reviewed, revised, and approved all content and take full responsibility for the published work.

## References

Abdi, H., & Valentin, D. 2007. "Multiple Correspondence Analysis." Encyclopedia of Measurement and Statistics.

Afanador, E., & Kjelland, M. 2025. "Determining Spatial Relationships between Landscape Structure and Avian Diversity in the Edwards Plateau of Texas." International Journal of Research Studies in Zoology, 9: 23–46. doi:10.20431/2454-941X.0901004

Aitken, S. N., Jordan, R., & Tumas, H. R. 2024. "Conserving Evolutionary Potential: Combining Landscape Genomics with Established Methods to Inform Plant Conservation." Annual Review of Plant Biology, 75: 707–736. doi:10.1146/annurev-arplant-070523-044239

Andrén, H. 1994. "Effects of Habitat Fragmentation on Birds and Mammals in Landscapes with Different Proportions of Suitable Habitat: A Review." Oikos, 71: 355–366. doi:10.2307/3545823

Arslan, D., Gaget, E., Çiçek, K., Olivier, A., Galewski, T., Döndüren, Ö., Guelmami, A., Ernoul, L., & Béchet, A. 2025. "Contrasting effects of agriculture and urbanisation on bird and reptile communities in a Mediterranean delta (Gediz Delta, Türkiye)." BMC Ecology and Evolution, 25: 58. doi:10.1186/s12862-025-02390-y

Asner, G. P., Archer, S., Hughes, R. F., Ansley, R. J., & Wessman, C. A. 2003. "Net changes in regional woody vegetation cover and carbon storage in Texas Drylands, 1937–1999." Global Change Biology, 9: 316–335. doi:10.1046/j.1365-2486.2003.00594.x

Bailey, V. 1905. Biological survey of Texas. North American Fauna No. 25. U.S. Government Printing Office.

Beirão, M. V., Neves, F. S., Penz, C. M., DeVries, P. J., & Fernandes, G. W. 2017. "High butterfly beta diversity between Brazilian cerrado and cerrado–caatinga transition zones." Journal of Insect Conservation, 21: 849–860. doi:10.1007/s10841-017-0024-x

Beja, P., & Alcazar, R. 2003. "Conservation of Mediterranean temporary ponds under agricultural intensification: an evaluation using amphibians." Biological Conservation, 114: 317–326. doi:10.1016/S0006-3207(03)00051-X

Beller, E. E., McClenachan, L., Zavaleta, E. S., & Larsen, L. G. 2020. "Past forward: Recommendations from historical ecology for ecosystem management." Global Ecology and Conservation, 21: e00836. doi:10.1016/j.gecco.2019.e00836

Benavidez, J. R., Dowell-Lashmet, T., Anderson, D. P., & Ullrich, K. 2021. "Converting Texas Land from Agricultural to Wildlife Use Tax Valuation." Journal of ASFMRA: 71–77.

Bevan, P. A., Ferreira, G. B., Ingram, D. J., Rowcliffe, M., Young, L., Freeman, R., & Jones, K. E. 2025. "Regional Biomes outperform broader spatial units in capturing biodiversity responses to land-use change." Ecography, 2025: e07318. doi:10.1111/ecog.07318

Biancolini, D., Riso, L., & Romano, A. 2024. "Opposite effects of climate and land use/cover change on Mediterranean herpetofauna: Insights from the southern Apennines." Animal Conservation, 28: 501–514. doi:10.1111/acv.12999

Bivand, R. (2025). spdep: Spatial dependence: weighting schemes, statistics (Version 1.4-1) [R Package]: CRAN.

Bosch, M. 2019. "PyLandStats: An open-source Pythonic library to compute landscape metrics." PLoS One. 14: e0225734. doi: 10.1371/journal.pone.0225734.

Breiman, L. 2001. "Random Forests." Machine Learning, 45: 5–32. doi:10.1023/A:1010933404324

Brooks, T., Mittermeier, R., Mittermeier, C., Fonseca, G., Rylands, A., Konstant, W., Flick, P., Pilgrim, J., Oldfield, S., Magin, G., & Hilton-Taylor, C. 2002. "Habitat Loss and Extinction in the Hotspots of Biodiversity." Conservation Biology, 16. doi:10.1046/j.1523-1739.2002.00530.x

Burnett, A. D., & Koprowski, J. L. 2024. "Shrub avoidance by an open-adapted ground squirrel in a shrub-encroached environment." PLoS One, 19: e0297993. doi:10.1371/journal.pone.0297993

Cain, M. D., & Shelton, M. G. 2001. "Secondary forest succession following reproduction cutting on the Upper Coastal Plain of southeastern Arkansas, USA." Forest Ecology and Management, 146: 223–238. doi:10.1016/S0378-1127(00)00464-3

Cartwright, J. M., & Wolfe, W. J. 2016. "Insular ecosystems of the southeastern United States—A regional synthesis to support biodiversity conservation in a changing climate." (1828). Retrieved from Nashville, TN.

Ceballos, G., Davidson, A., List, R., Pacheco, J., Manzano-Fischer, P., Santos-Barrera, G., & Cruzado, J. 2010. "Rapid Decline of a Grassland System and Its Ecological and Conservation Implications." PLoS One, 5: e8562. doi:10.1371/journal.pone.0008562

Ceballos, G., Ehrlich, P. R., & Dirzo, R. 2017. "Biological annihilation via the ongoing sixth mass extinction signaled by vertebrate population losses and declines." Proceedings of the National Academy of Sciences, 114: E6089–E6096. doi:doi:10.1073/pnas.1704949114

Chen, T.-t., Peng, L., Liu, S.-q., & Wang, Q. 2017. "Land cover change in different altitudes of Guizhou-Guangxi karst mountain area, China: patterns and drivers." Journal of Mountain Science, 14: 1873–1888. doi:10.1007/s11629-016-4202-1

Clavel, J., Julliard, R., & Devictor, V. 2011. "Worldwide decline of specialist species: toward a global functional homogenization?" Frontiers in Ecology and the Environment, 9: 222–228. doi:10.1890/080216

Colino-Rabanal, V., & Lizana, M. 2012. "Herpetofauna and roads: A review." Basic and Applied Herpetology, 26: 5–31. doi:10.11160/bah.12008

Comer, C. E., & Steumke, L. A. 2011. "Distribution and habitat requirements of bats in the Pineywoods ecoregion of east Texas, with emphasis on Rafinesque’s Big-eared Bat and Southeastern Myotis." Retrieved from Texas Parks and Wildlife Department.

Correll, M. D., Strasser, E. H., Green, A. W., & Panjabi, A. O. 2019. "Quantifying specialist avifaunal decline in grassland birds of the Northern Great Plains." Ecosphere, 10: e02523. doi:10.1002/ecs2.2523

Cox, N., Young, B. E., Bowles, P., Fernandez, M., Marin, J., Rapacciuolo, G., Böhm, M., Brooks, T. M., Hedges, S. B., Hilton-Taylor, C., Hoffmann, M., Jenkins, R. K. B., Tognelli, M. F., Alexander, G. J., Allison, A., Ananjeva, N. B., Auliya, M., Avila, L. J., Chapple, D. G., Cisneros-Heredia, D. F., Cogger, H. G., Colli, G. R., de Silva, A., Eisemberg, C. C., Els, J., Fong G, A., Grant, T. D., Hitchmough, R. A., Iskandar, D. T., Kidera, N., Martins, M., Meiri, S., Mitchell, N. J., Molur, S., Nogueira, C. d. C., Ortiz, J. C., Penner, J., Rhodin, A. G. J., Rivas, G. A., Rödel, M.-O., Roll, U., Sanders, K. L., Santos-Barrera, G., Shea, G. M., Spawls, S., Stuart, B. L., Tolley, K. A., Trape, J.-F., Vidal, M. A., Wagner, P., Wallace, B. P., & Xie, Y. 2022. "A global reptile assessment highlights shared conservation needs of tetrapods." Nature, 605: 285–290. doi:10.1038/s41586-022-04664-7

Cutler, D. R., Edwards, T. C., Jr., Beard, K. H., Cutler, A., Hess, K. T., Gibson, J., & Lawler, J. J. 2007. "Random forests for classification in ecology." Ecology, 88: 2783–2792. doi:10.1890/07-0539.1

Daskalova, G. N., Myers-Smith, I. H., & Godlee, J. L. 2020. "Rare and common vertebrates span a wide spectrum of population trends." Nature Communications, 11: 4394. doi:10.1038/s41467-020-17779-0

De Frenne, P., Graae, B. J., Rodríguez-Sánchez, F., Kolb, A., Chabrerie, O., Decocq, G., De Kort, H., De Schrijver, A., Diekmann, M., Eriksson, O., Gruwez, R., Hermy, M., Lenoir, J., Plue, J., Coomes, D. A., & Verheyen, K. 2013. "Latitudinal gradients as natural laboratories to infer species’ responses to temperature." Journal of Ecology, 101: 784–795. doi:10.1111/1365-2745.12074

Diamond, D. D., & Smiens, F. E. 1993. "The native plant communities of the Blackland Prairies." In Book "The native plant communities of the Blackland Prairies." (pp. 66–81). Waco, TX: Baylor University.

Diaz, S., Settele, J., Brondízio, E., Ngo, H. T., Agard, J., Arneth, A., Balvanera, P., Brauman, K., Butchart, S., Chan, K., Garibaldi, L., Ichii, K., Liu, J., Subramanian, S., Midgley, G., Miloslavich, P., Molnár, Z., Obura, D., Pfaff, A., & Zayas, C. 2019. "Pervasive human-driven decline of life on Earth points to the need for transformative change." Science (New York, N.Y.), 366. doi:10.1126/science.aax3100

Dodd, C. K. 2010. "Amphibian ecology and conservation: A handbook of techniques." Oxford, UK: Oxford University Press.

Donaldson, M. R., Burnett, N. J., Braun, D. C., Suski, C. D., Hinch, S. G., Cooke, S. J., & Kerr, J. T. 2016. "Taxonomic bias and international biodiversity conservation research." FACETS, 1: 105–113. doi:10.1139/facets-2016-0011

Doughty, R. W. 1986. "Settlement and Environmental Change in Texas, 1820-1900." The Southwestern Historical Quarterly, 89: 423–442.

Douglas, D. J. T., Waldinger, J., Buckmire, Z., Gibb, K., Medina, J. P., Sutcliffe, L., Beckmann, C., Collar, N. J., Jansen, R., Kamp, J., Little, I., Sheldon, R., Yanosky, A., & Koper, N. 2023. "A global review identifies agriculture as the main threat to declining grassland birds." Ibis, 165: 1107–1128. doi:10.1111/ibi.13223

Duren, O. C., Muir, P. S., & Hosten, P. E. 2012. "Vegetation Change from the Euro-American Settlement Era to the Present in Relation to Environment and Disturbance in Southwest Oregon." Northwest Science, 86: 310–328. doi:10.3955/046.086.0407

Esser, L. F., Neves, D., & Jarenkow, J. A. 2025. "Species Distribution Models to Help Integrate Community Ecology." Austral Ecology, 50: e70091. doi:10.1111/aec.70091

Fern, R. R., Baron, M. D., England, A. E., Giese, J. C., Kraai, K. J., Lancaster, J. D., Oldenburger, S. L., Shipes, J. C., Wilson, B. C., & Wyckoff, S. R. 2023. "The state of Texas wetlands: a review of current and future challenges." Texas Water Journal, 14.

Ferreira, M., & Beja, P. 2013. "Mediterranean amphibians and the loss of temporary ponds: Are there alternative breeding habitats?" Biological Conservation, 165: 179–186. doi:10.1016/j.biocon.2013.05.029

Foster, D., Swanson, F., Aber, J., Burke, I., Brokaw, N., Tilman, D., & Knapp, A. 2003. "The Importance of Land-Use Legacies to Ecology and Conservation." BioScience, 53: 77–88. doi:10.1641/0006-3568(2003)053[0077:TIOLUL]2.0.CO;2

Fukasawa, K., & Akasaka, T. 2019. "Long-lasting effects of historical land use on the current distribution of mammals revealed by ecological and archaeological patterns." Scientific Reports, 9: 10697. doi:10.1038/s41598-019-46809-1

García-Vega, D., & Newbold, T. 2020. "Assessing the effects of land use on biodiversity in the world’s drylands and Mediterranean environments." Biodiversity and Conservation, 29: 393–408. doi:10.1007/s10531-019-01888-4

Gaston, K. J. 2009. "Geographic range limits of species." Proceedings of the Royal Society B: Biological Sciences, 276: 1391–1393. doi:10.1098/rspb.2009.0100

Getis, A., & Ord, J. K. 1992. "The Analysis of Spatial Association by Use of Distance Statistics." Geographical Analysis, 24: 189–206. doi:10.1111/j.1538-4632.1992.tb00261.x

Griffith, G., Bryce, S., Omernik, J., & Rogers, A. 2007. "Ecoregions of Texas." Retrieved from Texas Commission on Environmental Quality.

Guo, J., & Zhang, M. 2021. "Exploring the Patterns and Drivers of Urban Expansion in the Texas Triangle Megaregion." Land, 10: 1244. doi:10.3390/land10111244

Hanberry, B. B. 2020. "Defining the Historical Northeastern Forested Boundary of the Great Plains Grasslands in the United States." The Professional Geographer, 72: 1–8. doi:10.1080/00330124.2019.1611460

Hayes, T. B., Falso, P., Gallipeau, S., & Stice, M. 2010. "The cause of global amphibian declines: a developmental endocrinologist’s perspective." Journal of Experimental Biology, 213: 921–933. doi:10.1242/jeb.040865

Holt, E. A., Allen, K. E., Parker, N. C., Baker, R. J. 2000. "Ecotourism and conservation: Richness of terrestrial vertebrates across Texas." Museum of Texas Tech University Occasional Paper OP-201.16p

Hu, J., Long, Y., Zhou, W., Zhu, C., Yang, Q., Zhou, S., & Wu, P. 2020. "Influence of different land use types on hydrochemistry and heavy metals in surface water in the lakeshore zone of the Caohai wetland, China." Environmental Pollution, 267: 115454. doi:10.1016/j.envpol.2020.115454

Johnston, C. A. 2014. "Agricultural expansion: land use shell game in the U.S. Northern Plains." Landscape Ecology, 29: 81–95. doi:10.1007/s10980-013-9947-0

Jung, M., Dahal, P. R., Butchart, S. H. M., Donald, P. F., De Lamo, X., Lesiv, M., Kapos, V., Rondinini, C., & Visconti, P. 2020. "A global map of terrestrial habitat types." Scientific Data, 7: 256. doi:10.1038/s41597-020-00599-8

Kéfi, S., Génin, A., Garcia-Mayor, A., Guirado, E., Cabral, J. S., Berdugo, M., Guerber, J., Solé, R., & Maestre, F. T. 2024. "Self-organization as a mechanism of resilience in dryland ecosystems." Proceedings of the National Academy of Sciences, 121: e2305153121. doi:doi:10.1073/pnas.2305153121

Kosti, M., Lazaridou, S., Bourazani, N., & Angelis, L. 2011. "Discovering patterns of correlation and similarities in software project data with the Circos visualization tool."

Kreuter, U. P., Wolfe, D. W., Hays, K. B., & Conner, J. R. 2017. "Conservation Credits—Evolution of a Market-Oriented Approach to Recovery of Species of Concern on Private Land." Rangeland Ecology & Management, 70: 264–272. doi:10.1016/j.rama.2016.10.012

Krzywinski, M., Schein, J., Birol, I., Connors, J., Gascoyne, R., Horsman, D., Jones, S. J., & Marra, M. A. 2009. "Circos: an information aesthetic for comparative genomics." Genome Research, 19: 1639–1645. doi:10.1101/gr.092759.109

Kuykendall, M. T., & Gregory, S. K. 2011. "Impacts of Roads and Corridors on Abundance and Movement of Small Mammals on the Llano Estacado of Texas." The Southwestern Naturalist, 56: 9–16. doi:10.1894/CLG-33.1

Labay, B., Cohen, A. E., Sissel, B., Hendrickson, D. A., Martin, F. D., & Sarkar, S. 2011. "Assessing historical fish community composition using surveys, historical collection data, and species distribution models." PLoS One, 6: e25145. doi:10.1371/journal.pone.0025145

Lê, S., Josse, J., & Husson, F. 2008. "FactoMineR: An R Package for Multivariate Analysis." Journal of Statistical Software, 25: 1 – 18. doi:10.18637/jss.v025.i01

Li, Q., Shao, W., Jiang, Y., Yan, C., & Liao, W. 2024. "Assessing Reptile Conservation Status under Global Climate Change." Biology, 13: 436. doi:10.3390/biology13060436

Li, X., Tian, H., Lu, C., & Pan, S. 2023. "Four-century history of land transformation by humans in the United States (1630–2020): annual and 1 km grid data for the history of land changes (HISLAND-US)." Earth System Science Data, 15: 1005–1035. doi:10.5194/essd-15-1005-2023

Lindenmayer, D., Scheele, B. C., Lavery, T., & Likens, G. E. 2023. "Biodiversity response to rapid successive land cover conversions in human-dominated landscapes." Global Ecology and Conservation, 45: e02510. doi:10.1016/j.gecco.2023.e02510

Lombardi, J., Perotto-Baldivieso, H., & Tewes, M. 2020. "Land Cover Trends in South Texas (1987–2050): Potential Implications for Wild Felids." Remote Sensing, 12: 659. doi:10.3390/rs12040659

Luedtke, J. A., Chanson, J., Neam, K., Hobin, L., Maciel, A. O., Catenazzi, A., Borzée, A., Hamidy, A., Aowphol, A., Jean, A., Sosa-Bartuano, Á., Fong G, A., de Silva, A., Fouquet, A., Angulo, A., Kidov, A. A., Muñoz Saravia, A., Diesmos, A. C., Tominaga, A., Shrestha, B., Gratwicke, B., Tjaturadi, B., Martínez Rivera, C. C., Vásquez Almazán, C. R., Señaris, C., Chandramouli, S. R., Strüssmann, C., Cortez Fernández, C. F., Azat, C., Hoskin, C. J., Hilton-Taylor, C., Whyte, D. L., Gower, D. J., Olson, D. H., Cisneros-Heredia, D. F., Santana, D. J., Nagombi, E., Najafi-Majd, E., Quah, E. S. H., Bolaños, F., Xie, F., Brusquetti, F., Álvarez, F. S., Andreone, F., Glaw, F., Castañeda, F. E., Kraus, F., Parra-Olea, G., Chaves, G., Medina-Rangel, G. F., González-Durán, G., Ortega-Andrade, H. M., Machado, I. F., Das, I., Dias, I. R., Urbina-Cardona, J. N., Crnobrnja-Isailović, J., Yang, J.-H., Jianping, J., Wangyal, J. T., Rowley, J. J. L., Measey, J., Vasudevan, K., Chan, K. O., Gururaja, K. V., Ovaska, K., Warr, L. C., Canseco-Márquez, L., Toledo, L. F., Díaz, L. M., Khan, M. M. H., Meegaskumbura, M., Acevedo, M. E., Napoli, M. F., Ponce, M. A., Vaira, M., Lampo, M., Yánez-Muñoz, M. H., Scherz, M. D., Rödel, M.-O., Matsui, M., Fildor, M., Kusrini, M. D., Ahmed, M. F., Rais, M., Kouamé, N. G. G., García, N., Gonwouo, N. L., Burrowes, P. A., Imbun, P. Y., Wagner, P., Kok, P. J. R., Joglar, R. L., Auguste, R. J., Brandão, R. A., Ibáñez, R., von May, R., Hedges, S. B., Biju, S. D., Ganesh, S. R., Wren, S., Das, S., Flechas, S. V., Ashpole, S. L., Robleto-Hernández, S. J., Loader, S. P., Incháustegui, S. J., Garg, S., Phimmachak, S., Richards, S. J., Slimani, T., Osborne-Naikatini, T., Abreu-Jardim, T. P. F., Condez, T. H., De Carvalho, T. R., Cutajar, T. P., Pierson, T. W., Nguyen, T. Q., Kaya, U., Yuan, Z., Long, B., Langhammer, P., & Stuart, S. N. 2023. "Ongoing declines for the world’s amphibians in the face of emerging threats." Nature, 622: 308–314. doi:10.1038/s41586-023-06578-4

Mallet, P., Bechet, A., Sirami, C., Mesleard, F., Blanchon, T., Calatayud, F., Dagonet, T., Gaget, E., Leray, C., & Galewski, T. 2023. "Field margins as substitute habitat for the conservation of birds in agricultural wetlands." Peer Community Journal, 3. doi:10.24072/pcjournal.299

Martins, I. S., Dornelas, M., Vellend, M., & Thomas, C. D. 2022. "A millennium of increasing diversity of ecosystems until the mid-20th century." Global Change Biology, 28: 5945–5955. doi:10.1111/gcb.16335

McDonald, R., McKnight, M., Weiss, D., Selig, E., O’Connor, M., Violin, C., & Moody, A. 2005. "Species compositional similarity and ecoregions: Do ecoregion boundaries represent zones of high species turnover?" Biological Conservation, 126: 24–40. doi:10.1016/j.biocon.2005.05.008

Meineke, E. K., Davies, T. J., Daru, B. H., & Davis, C. C. 2018. "Biological collections for understanding biodiversity in the Anthropocene." Philosophical Transactions of the Royal Society B: Biological Sciences, 374: 20170386. doi:10.1098/rstb.2017.0386

Meyer, D., Zeileis, A., Hornik, K., & Friendly, M. (2015). vcd: Visualizing categorical data (Version R package version 1.4-13): CRAN.

Mi, C., Huettmann, F., Guo, Y., Han, X., & Wen, L. 2017. "Why choose Random Forest to predict rare species distribution with few samples in large undersampled areas? Three Asian crane species models provide supporting evidence." PeerJ, 5: e2849. doi:10.7717/peerj.2849

Mullu, D. 2016. "A Review on the Effect of Habitat Fragmentation on Ecosystem." Journal of Natural Sciences Research, 6.

Murphy, M. O., Farleigh, K., Peterman, W. E., Jezkova, T., & Boone, M. D. 2025. "Land-cover patterns differentially affect population genetic structure and connectivity of two anurans." Landscape Ecology, 40: 74. doi:10.1007/s10980-025-02086-0

Newbold, T., Adams, G. L., Albaladejo Robles, G., Boakes, E. H., Braga Ferreira, G., Chapman, A. S. A., Etard, A., Gibb, R., Millard, J., Outhwaite, C. L., & Williams, J. J. 2019. "Climate and land-use change homogenise terrestrial biodiversity, with consequences for ecosystem functioning and human well-being." Emerging Topics in Life Sciences, 3: 207–219. doi:10.1042/ETLS20180135

North American Bird Conservation Initiative. 2025. "The state of the birds, United States of America, 2025." Retrieved from Stateofthebirds.org

O’Reilly, E., Gregory, R. D., Aunins, A., Brotons, L., Chodkiewicz, T., Escandell, V., Foppen, R. P. B., Gamero, A., Herrando, S., Jiguet, F., Kålås, J. A., Kamp, J., Klvaňová, A., Lehikoinen, A., Lindström, Å., Massimino, D., Jostein Øien, I., Reif, J., Šilarová, E., Teufelbauer, N., Trautmann, S., van Turnhout, C., Vikstrøm, T., Voříšek, P., & Butler, S. J. 2022. "An assessment of relative habitat use as a metric for species’ habitat association and degree of specialization." Ecological Indicators, 135: 108521. 10.1016/j.ecolind.2021.108521

Olariu, H., Wilcox, B., & Popescu, S. 2024. "Examining changes in woody vegetation cover in a human-modified temperate savanna in Central Texas between 1996 and 2022 using remote sensing." Frontiers in Forests and Global Change, 7. doi:10.3389/ffgc.2024.1396999

Oliver, R. Y., Meyer, C., Ranipeta, A., Winner, K., & Jetz, W. 2021. "Global and national trends, gaps, and opportunities in documenting and monitoring species distributions." PLoS Biology, 19: e3001336. doi:10.1371/journal.pbio.3001336

Pacifici, M., Visconti, P., Butchart, S. H. M., Watson, J. E. M., Cassola, Francesca M., & Rondinini, C. 2017. "Species’ traits influenced their response to recent climate change." Nature Climate Change, 7: 205–208. doi:10.1038/nclimate3223

Pereira, A. A., Rosa, C., Faria, L. D. B., Silva, L. G. D., & Passamani, M. 2023. "Human presence as a determinant of the occurrence of mammals in a high diversity protected area of Cerrado-Caatinga ecotone in Brazil." Anais da Academia Brasileira de Ciencias, 95: e20201869. doi:10.1590/0001-3765202320201869

Pimm, S. L., Jenkins, C. N., Abell, R., Brooks, T. M., Gittleman, J. L., Joppa, L. N., Raven, P. H., Roberts, C. M., & Sexton, J. O. 2014. "The biodiversity of species and their rates of extinction, distribution, and protection." Science, 344: 1246752. doi:doi:10.1126/science.1246752

Provost, G. L., Badenhausser, I., Le Bagousse-Pinguet, Y., Clough, Y., Henckel, L., Violle, C., Bretagnolle, V., Roncoroni, M., Manning, P., & Gross, N. 2020. "Land-use history impacts functional diversity across multiple trophic groups." Proceedings of the National Academy of Sciences, 117: 1573–1579. doi:doi:10.1073/pnas.1910023117

Pyne, S. J. 1997. "America’s fires: Management on wildlands and forests." Seattle: University of Washington Press.

Quintas-Soriano, C., Buerkert, A., & Plieninger, T. 2022. "Effects of land abandonment on nature contributions to people and good quality of life components in the Mediterranean region: A review." Land Use Policy, 116: 106053. doi:10.1016/j.landusepol.2022.106053

Ren, B., Park, K., Shrestha, A., Yang, J., McHale, M., Bai, W., & Wang, G. 2022. "Impact of Human Disturbances on the Spatial Heterogeneity of Landscape Fragmentation in Qilian Mountain National Park, China." Land, 11: 2087. doi:10.3390/land11112087

Rideout, D. W. 1994. "The Post Oak Savannah deer herd: Past, present, and future." Retrieved from College Station, TX.

Ritchie, A. 2025. "Paved paradise: Using property tax incentives to curb urban sprawl in Texas communities." SSRN.

Robertson, H., Fennessy, S., Hilton, G., Job, N., Kumar, R., Peled, Y., Simpson, M., Aggestam, F., Eldred, M., Davidson, N., Costanza, R., Chacon-Cascante, A., Field, C., Finlayson, M., Gandra, F., Gillis, L., Hernández-Blanco, M., Moritsch, M., Thornton, S., & Hoff, V. 2025. "Global Wetland Outlook 2025: Valuing, conserving, restoring and financing wetlands."

Roos, C. I., Zedeño, M. N., Hollenback, K. L., & Erlick, M. M. H. 2018. "Indigenous impacts on North American Great Plains fire regimes of the past millennium." Proceedings of the National Academy of Sciences, 115: 8143–8148. doi:doi:10.1073/pnas.1805259115

Rosenberg, K. V., Dokter, A. M., Blancher, P. J., Sauer, J. R., Smith, A. C., Smith, P. A., Stanton, J. C., Panjabi, A., Helft, L., Parr, M., & Marra, P. P. 2019. "Decline of the North American avifauna." Science, 366: 120–124. doi:10.1126/science.aaw1313

Schmandt, J., North, G. R., & Clarkson, J. 2011. "The Impact of Global Warming on Texas*: Second edition."* University of Texas Press.

Schmidley, D. J., Bradley, R. D., Bradley, L. C. 2022. “Texas Natural History in the 21st Century.” Lubbock TX: Texas Tech University Press.

Schmitt, C. J., Cook, J. A., Zamudio, K. R., & Edwards, S. V. 2018. "Museum specimens of terrestrial vertebrates are sensitive indicators of environmental change in the Anthropocene." Philosophical Transactions of the Royal Society B: Biological Sciences, 374: 20170387. doi:10.1098/rstb.2017.0387

Simon, S. M., Glaum, P., & Valdovinos, F. S. 2023. "Interpreting random forest analysis of ecological models to move from prediction to explanation." Scientific Reports, 13: 3881. doi:10.1038/s41598-023-30313-8

Smith, A., Lopez, R., Lund, A., Anderson, R., Wegner, B., Anderson, A., Crawford, M., Powers, G., Skow, K., & Lopez, A. 2025. "*Status update and trends of Texas working lands."* Retrieved from College Station, TX.

Smith, J. R., Letten, A. D., Ke, P. J., Anderson, C. B., Hendershot, J. N., Dhami, M. K., Dlott, G. A., Grainger, T. N., Howard, M. E., Morrison, B. M. L., Routh, D., San Juan, P. A., Mooney, H. A., Mordecai, E. A., Crowther, T. W., & Daily, G. C. 2018. "A global test of ecoregions." Nature Ecology & Evolution, 2: 1889–1896. doi:10.1038/s41559-018-0709-x

Sohl, T. L., Reker, R., Bouchard, M., Sayler, K., Dornierer, J., Wika, S., Quenzer, R., & Friesz, A. 2018. "Modeled historical land use and land cover for the conterminous United States: 1938-1992." U.S. Geological Survey. doi:10.5066/F7KK99RR

Solem, E., Radchuk, V., Schwarz, J., Kramer-Schadt, S., & Planillo, A. 2025. "Multi-scale analysis reveals vegetation buffers human disturbance impacts on urban bird functional diversity." Urban Ecosystems, 28: 223. doi:10.1007/s11252-025-01829-w

Strobl, C., Boulesteix, A.-L., Kneib, T., Augustin, T., & Zeileis, A. 2008. "Conditional variable importance for random forests." BMC Bioinformatics, 9: 307. doi:10.1186/1471-2105-9-307

Suarez, A. V., & Tsutsui, N. D. 2004. "The Value of Museum Collections for Research and Society." BioScience, 54: 66–74. doi:10.1641/0006-3568(2004)054[0066:TVOMCF]2.0.CO;2

Subba, B., Sen, S., Ravikanth, G., & Nobis, M. P. 2018. "Direct modelling of limited migration improves projected distributions of Himalayan amphibians under climate change." Biological Conservation, 227: 352–360. doi:10.1016/j.biocon.2018.09.035

Surasinghe, T., & Baldwin, R. F. 2014. "Ghost of land-use past in the context of current land cover: evidence from salamander communities in streams of Blue Ridge and Piedmont ecoregions." Canadian Journal of Zoology, 92: 527–536. doi:10.1139/cjz-2013-0307

Swetnam, T. W., Farella, J., Roos, C. I., Liebmann, M. J., Falk, D. A., & Allen, C. D. 2016. "Multiscale perspectives of fire, climate and humans in western North America and the Jemez Mountains, USA." Philosophical Transactions of the Royal Society B: Biological Sciences, 371. doi:10.1098/rstb.2015.0168

Tewksbury, J. J., Levey, D. J., Haddad, N. M., Sargent, S., Orrock, J. L., Weldon, A., Danielson, B. J., Brinkerhoff, J., Damschen, E. I., & Townsend, P. 2002. "Corridors affect plants, animals, and their interactions in fragmented landscapes." Proceedings of the National Academy of Sciences, 99: 12923–12926. doi:10.1073/pnas.202242699

Texas A&M Forest Service. 2009. "Texas statewide assessment of forest resources." Retrieved from College Station, TX.

Thorne, J., Choe, H., Dorji, L., Yangden, K., Wangdi, D., Phuntsho, Y., & Beardsley, K. 2022. "Species richness and turnover patterns for tropical and temperate plants on the elevation gradient of the eastern Himalayan Mountains." Frontiers in Ecology and Evolution, 10: 942759. doi:10.3389/fevo.2022.942759

Titley, M. A., Snaddon, J. L., & Turner, E. C. 2017. "Scientific research on animal biodiversity is systematically biased towards vertebrates and temperate regions." PLoS One, 12: e0189577. doi:10.1371/journal.pone.0189577

Todd, B. D., & Rothermel, B. B. 2006. "Assessing quality of clearcut habitats for amphibians: Effects on abundances versus vital rates in the southern toad (Bufo terrestris)." Biological Conservation, 133: 178–185. doi:10.1016/j.biocon.2006.06.003

Torre, I., & Palau, O. 2023. "Is Shrub Encroachment Driving the Decline of Small Mammal Diversity in Pyrenean Grasslands? A Preliminary Study." Diversity, 15: 232.

TPWD. 2002. "Land and water resources conservation and recreation plan." Retrieved from Austin, TX.

TPWD. 2006. "Texas wildlife action plan." Retrieved from Austin, TX.

TPWD. 2020. "Trans-Pecos Ecoregion." Retrieved from https://tpwd.texas.gov/wildlife/wildlife-diversity/wildscapes/wildscapes-plant-guidance-by-ecoregion/the-trans-pecos/

Turner, M. G. 2001. "Landscape ecology in theory and practice: pattern and process." (2 ed.). New York City, NY: Springer.

U.S. Geological Survey. 2024. "Annual NLCD collection 1 science products (ver. 1.1, June 2025)." U.S. Geological Survey. doi:10.5066/P94UXNTS

Vajana, E., Bozzano, M., Marchi, M., & Piotti, A. 2022. "On the Inclusion of Adaptive Potential in Species Distribution Models: Towards a Genomic-Informed Approach to Forest Management and Conservation." Environments, 10: 3. doi:10.3390/environments10010003

Vargas-Jaimes, J., González-Fernández, A., Joaquín Torres-Romero, E., Bolom-Huet, R., Manjarrez, J., Gopar-Merino, F., P. Pacheco, X., Garrido-Garduño, T., Chávez, C., & Sunny, A. 2021. "Impact of climate and land cover changes on the potential distribution of four endemic salamanders in Mexico." Journal for Nature Conservation, 64: 126066. doi:10.1016/j.jnc.2021.126066

Veselka, A., Aponte-Gutiérrez, A., Medina-Báez, O., & Watling, J. 2023. "Upper thermal limits predict herpetofaunal responses to forest edge and cover." Biotropica, 55. doi:10.1111/btp.13208

Wall, T. P., Oswald, B. P., Kidd, K. R., & Darville, R. L. 2019. "An evaluation of United States forest Service prescribed fire regimes in East Texas." Forest Ecology and Management, 449: 117485. doi:10.1016/j.foreco.2019.117485

Wambulwa, M. C., Milne, R., Wu, Z. Y., Spicer, R. A., Provan, J., Luo, Y. H., Zhu, G. F., Wang, W. T., Wang, H., Gao, L. M., Li, D. Z., & Liu, J. 2021. "Spatiotemporal maintenance of flora in the Himalaya biodiversity hotspot: Current knowledge and future perspectives." Ecol Evol, 11: 10794–10812. doi:10.1002/ece3.7906

Wang, L., Jiao, W., MacBean, N., et al. 2022. Dryland productivity under a changing climate. Nature Climate Change, 12: 981–994. doi:10.1038/s41558-022-01499-y

Webb, W. L. 1950. "Biogeographic Regions of Texas and Oklahoma." Ecology, 31: 426–433. doi:10.2307/1931496

Werneck, F. P. 2011. "The diversification of eastern South American open vegetation biomes: Historical biogeography and perspectives." Quaternary Science Reviews, 30: 1630–1648. doi:10.1016/j.quascirev.2011.03.009

Westaway, D. M., Nimmo, D. G., Jolly, C. J., Michael, D. R., Watson, D. M., & von Takach, B. 2025. "Genomic repercussions of landscape modification on three lizard species." Conservation Genetics, 26: 529–543. doi:10.1007/s10592-025-01686-2

Wilcox, B. P., & Huang, Y. 2010. "Woody plant encroachment paradox: Rivers rebound as degraded grasslands convert to woodlands." Geophysical Research Letters, 37. doi:10.1029/2009GL041929

Wilcox, B. P., Sorice, M. G., Angerer, J., & Wright, C. L. 2012. "Historical Changes in Stocking Densities on Texas Rangelands." Rangeland Ecology & Management, 65: 313–317. doi:10.2111/REM-D-11-00119.1

Wilkins, N. R., Snelgrove, A. G., Fitzsimons, B. C., Stevener, B. M., Skow, K. L., Anderson, R. E., & Dube, A. M. 2009. "Current land use trends (Texas land trends)." Texas A&M Institute of Renewable Natural Resources.

Williams, J. J., Freeman, R., Spooner, F., & Newbold, T. 2022. "Vertebrate population trends are influenced by interactions between land use, climatic position, habitat loss and climate change." Glob Chang Biol, 28: 797–815. doi:10.1111/gcb.15978

WWF. 2018. "Living planet report - 2018. Aiming higher." (978-2-940529-90-2). Retrieved from Gland, Switzerland.

Yancey, F. D., Schmidley, D. J., Kasper, S., & Manning, R. W. 2023. "The mammals of Trans-Pecos Texas: Including Big Bend and Guadalupe Mountains National Parks." College Station, TX: Texas A&M University Press.

