## Supplemental Information for "Legacy Effects of Land-Use Land-Cover Change on Species Distribution Dynamics: Evidence from a landscape of diverse biogeographic crossroads"

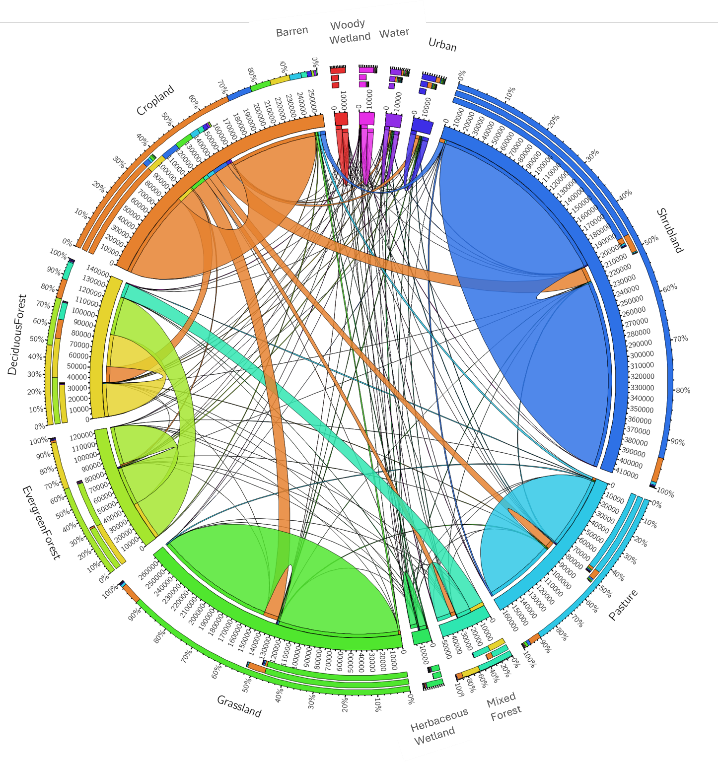

**Figure S1.** Chord diagrams of land-use and land-cover transitions between 1938-1992.

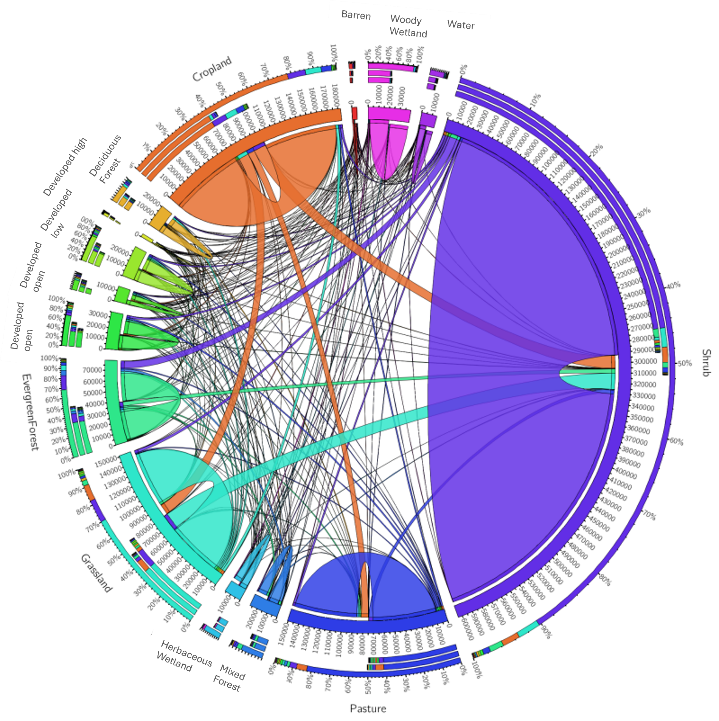

**Figure S2.** Chord diagrams of land-use and land-cover transitions between 1985-2023.

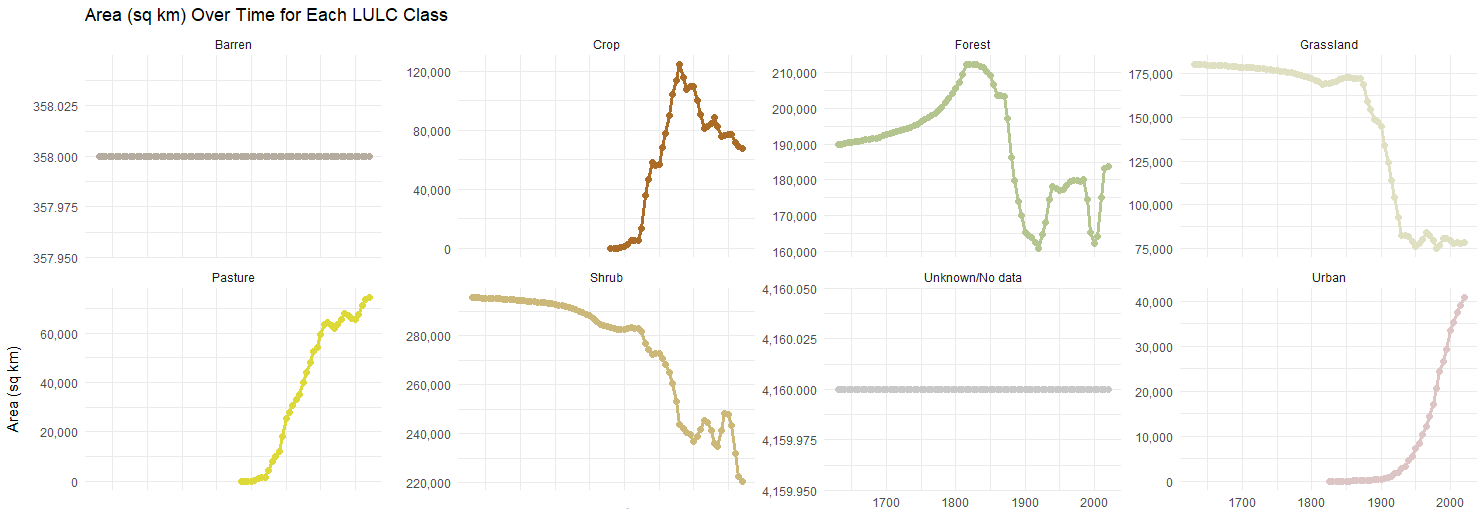

1700	1800	1900	2000

1700	1800	1900	2000

1700	1800	1900	2000

**Figure S3.** Percent land cover trends in Texas by class from 1850-2020.

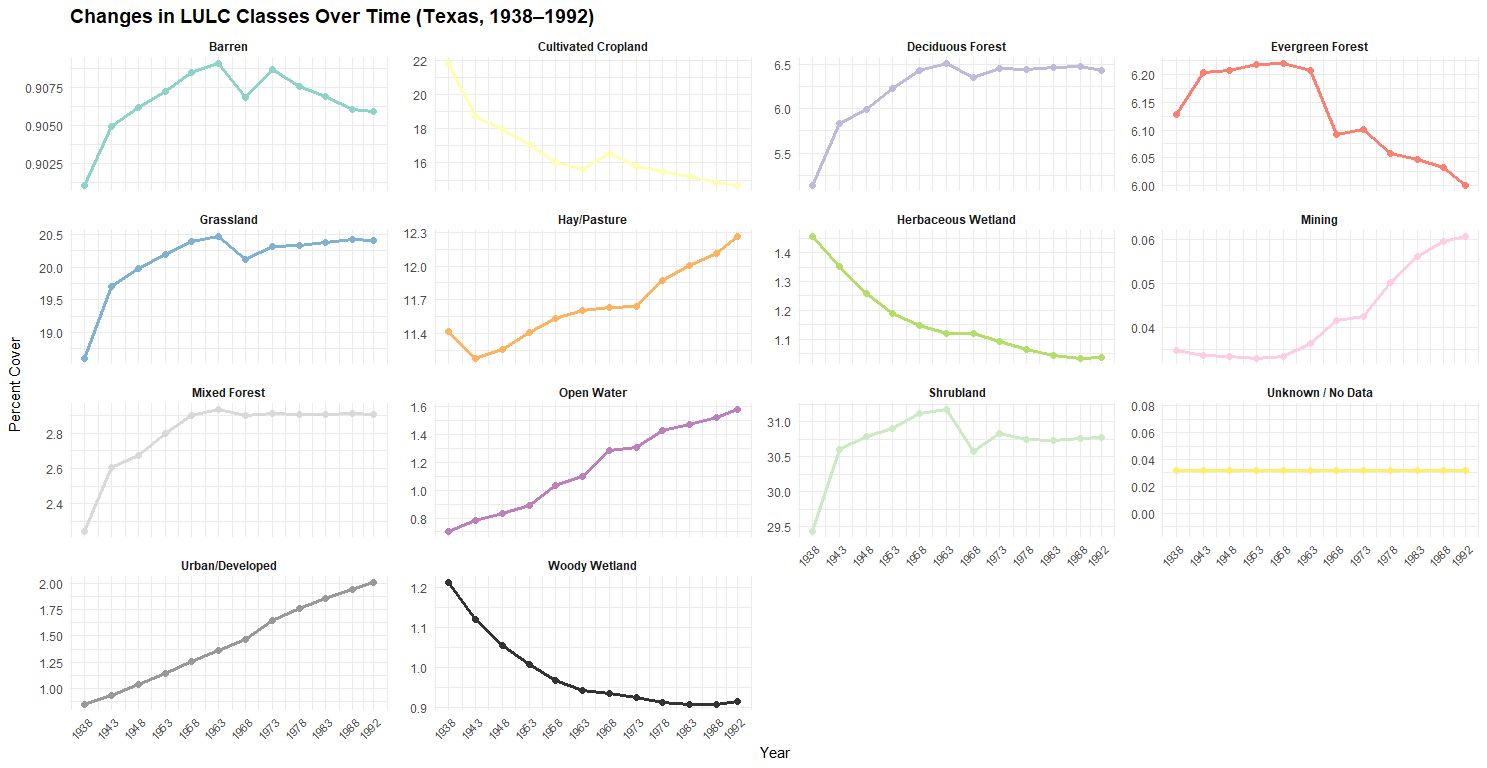

**Figure S4.** Percent land cover trends in Texas by class from 1938-1992.

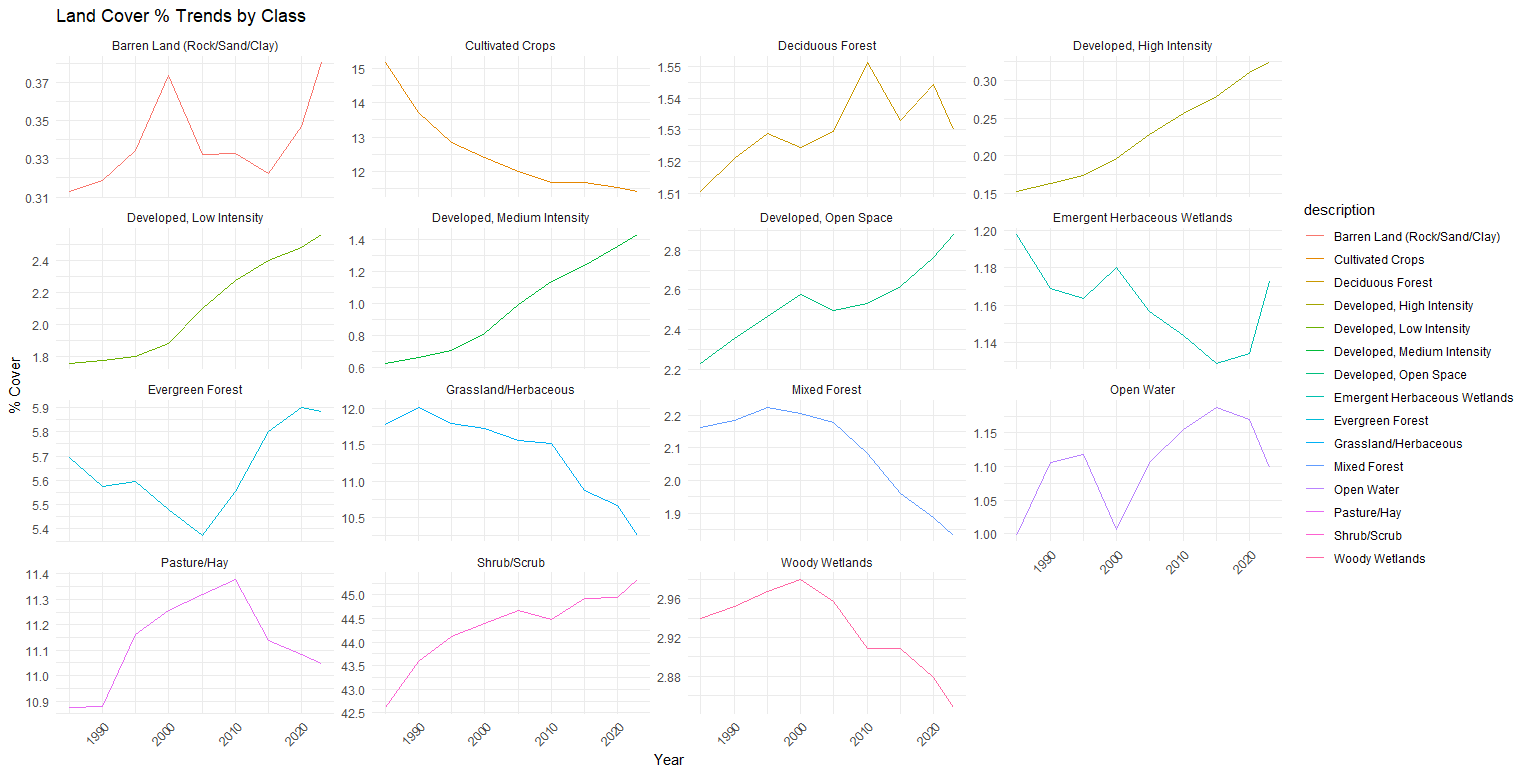

**Figure S5.** Percent land cover trends in Texas by class from 1985-2023.

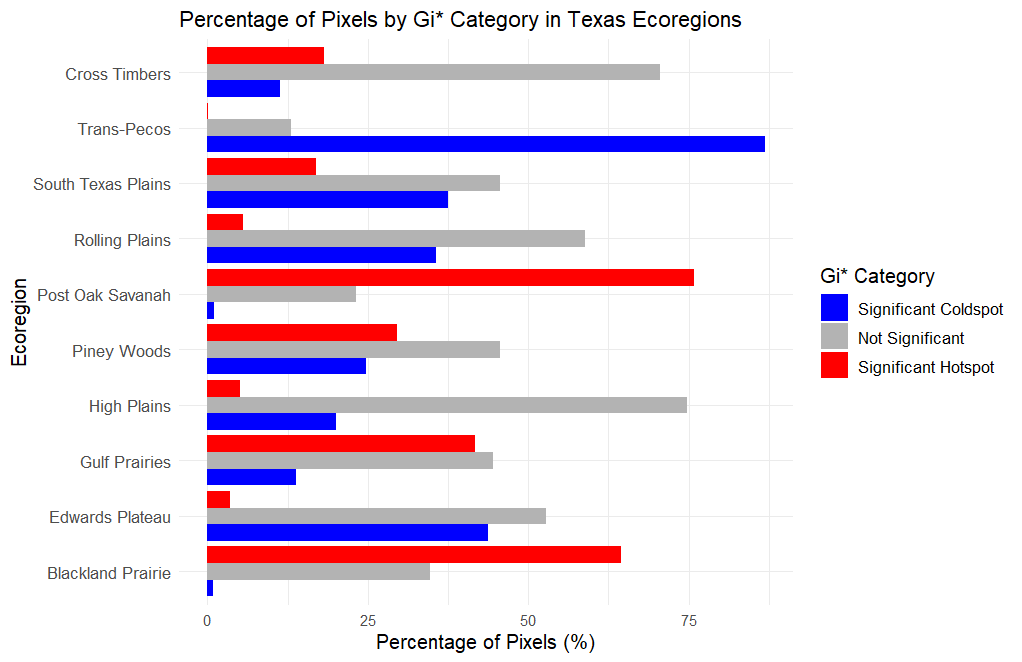

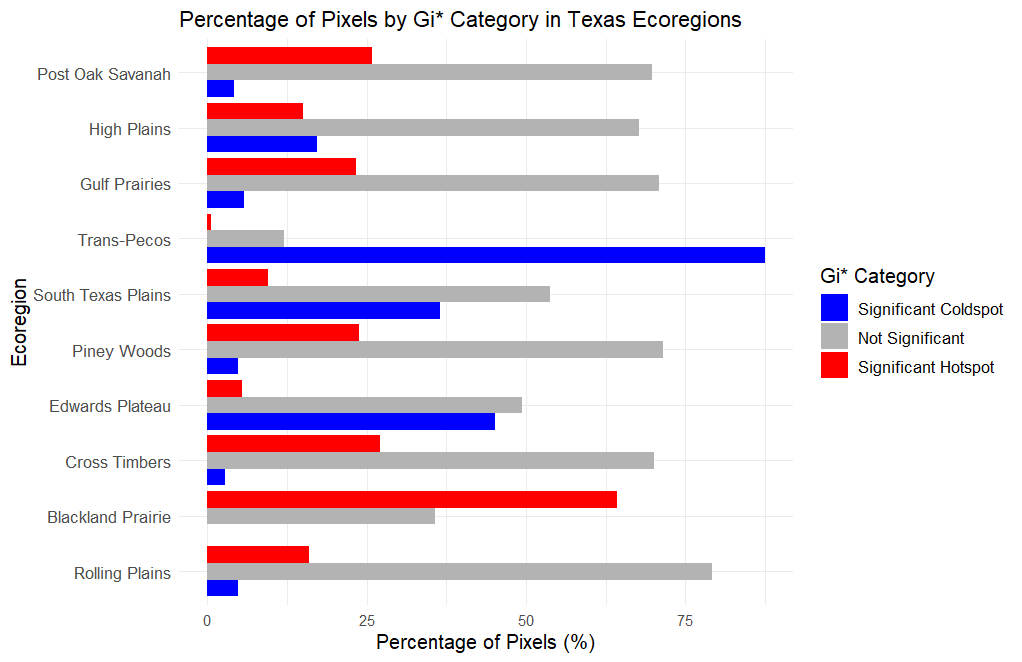

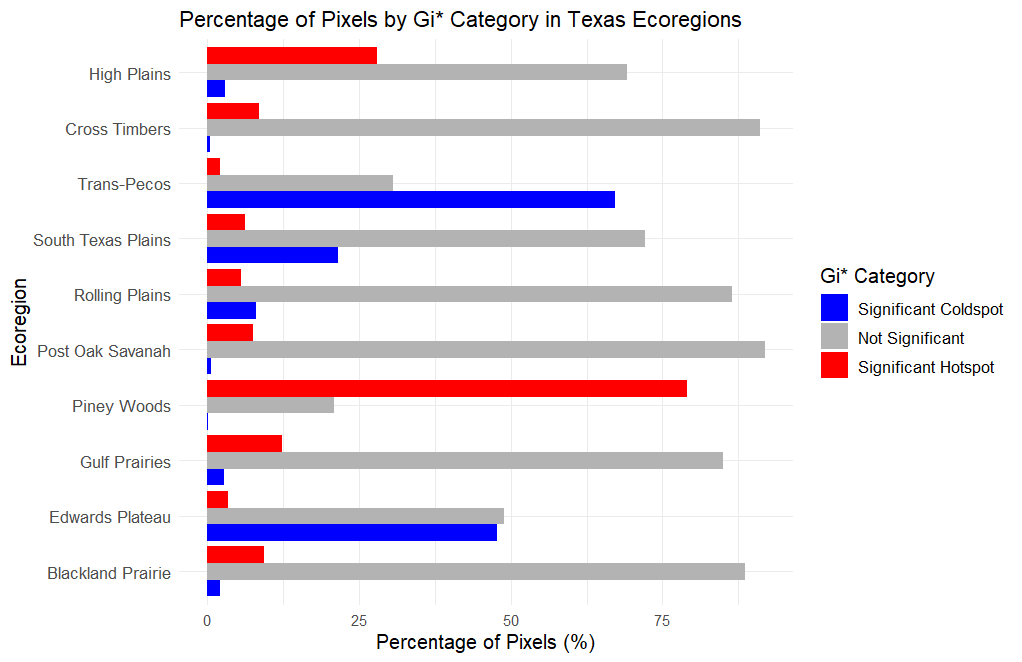

C)

A)

B)

**Figure S6.** Percentage of pixels (as 2.5 km x 2.5 km) classified by Getis-Ord Gi* statistic as hotspot categories within Texas ecoregions for land cover change from A) 1850-2020, B) 1938-1992, and C) 1985-2023. Red columns indicate statistically significant hotspots of land cover change, where high magnitudes of change were observed in surrounding areas. Blue columns indicate statistically significant coldspots, where there was very little magnitude of change observed in surrounding areas. Grey columns indicate locations that did not have significant land cover change.

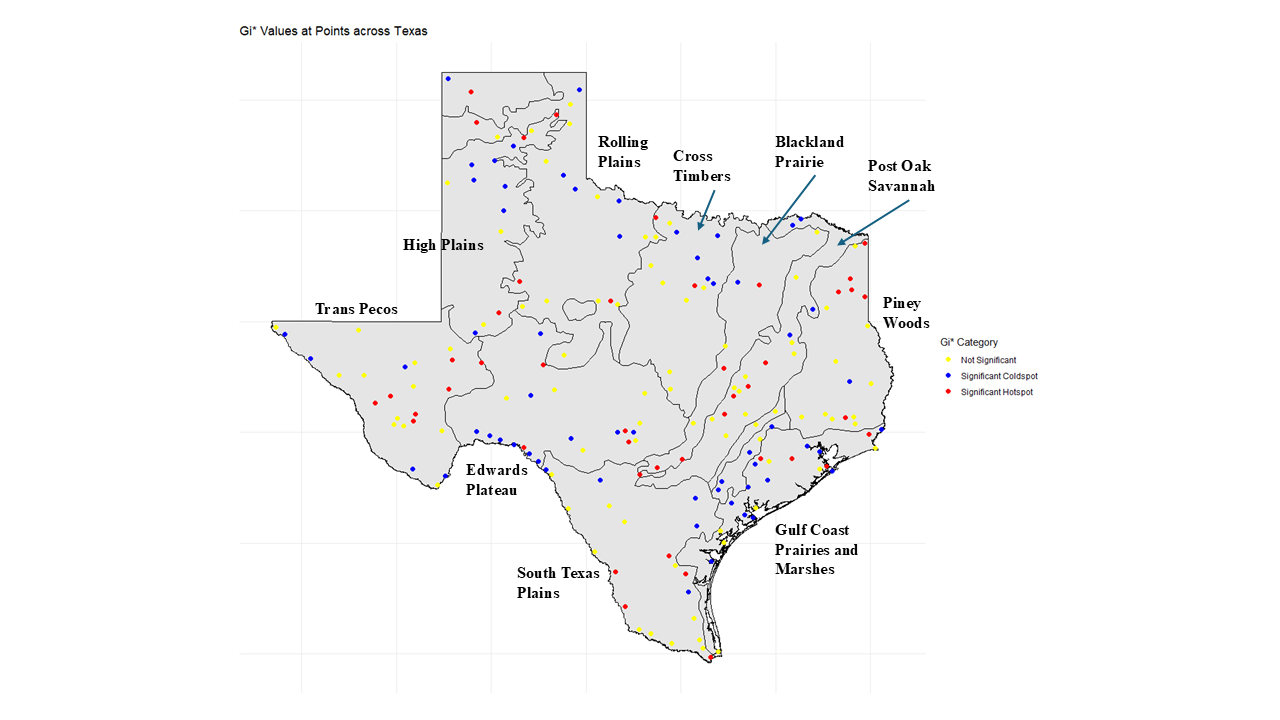

**Figure S7.** Getis-Ord Gi* hotspot analysis of land cover change across Texas between 1850-2020 using Bailey’s original survey points with outlines of ecoregion boundaries. Points are classified as statistically significant hotspots (red), coldspots (blue), or not significant (yellow) based on local clustering of land cover change.

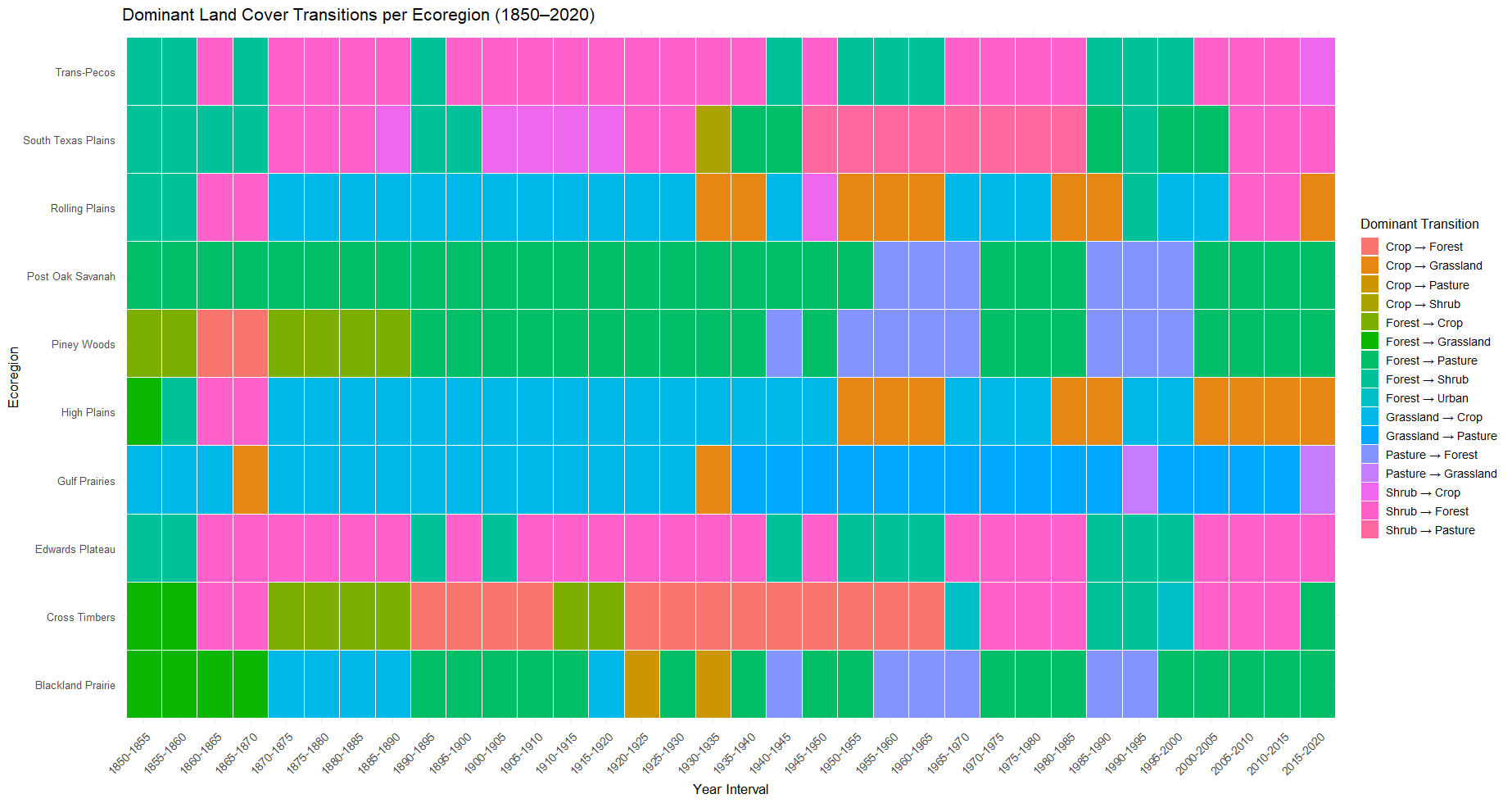

**Figure S8.** Dominant land cover transitions in each ecoregion of Texas from 1850-2020. The heatmap summarizes the most frequent land cover transition observed within each ecoregion during 5-year intervals.

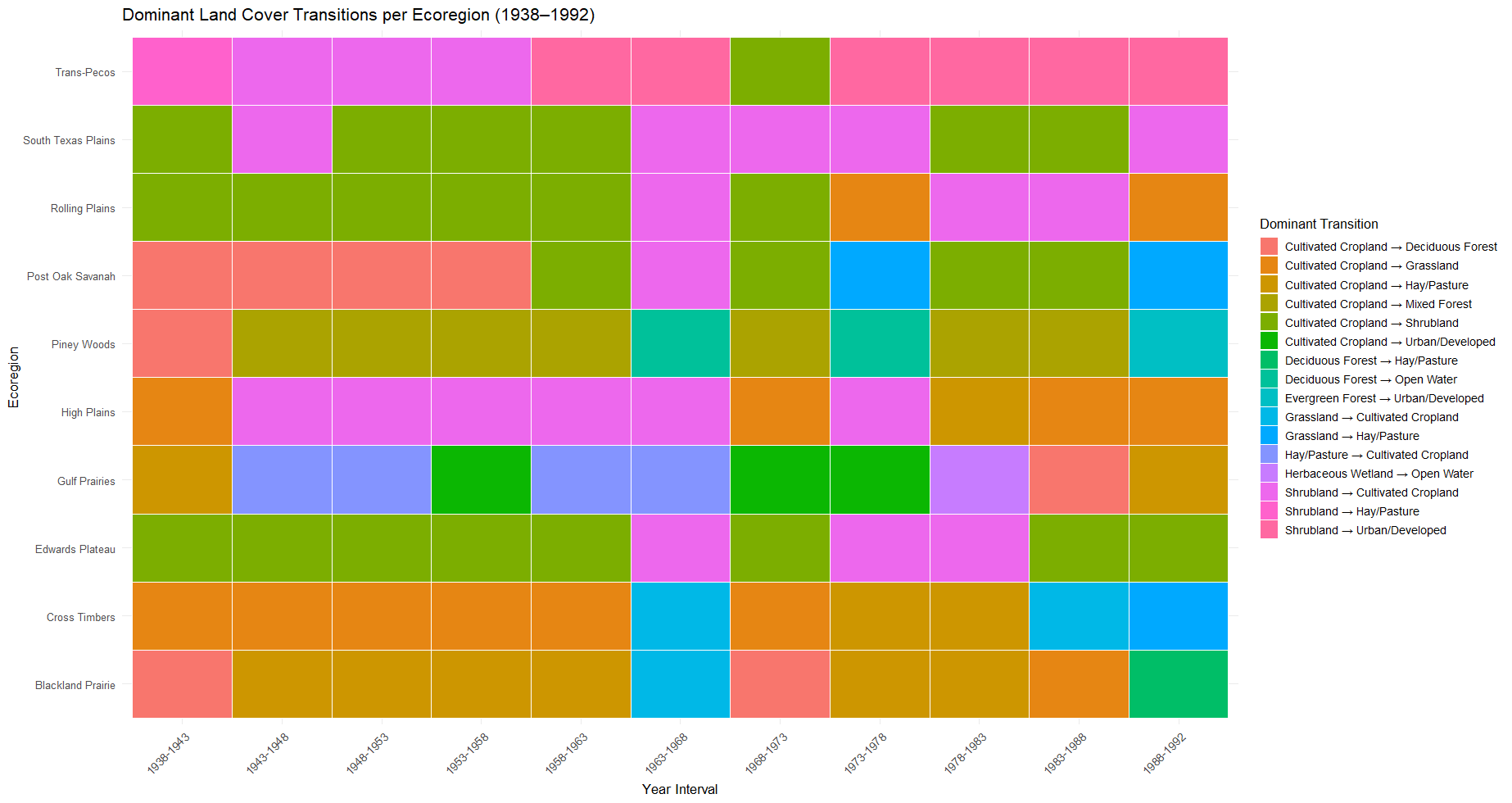

**Figure S9.** Dominant land cover transitions in each ecoregion of Texas from 1938-1992. The heatmap summarizes the most frequent land cover transition observed within each ecoregion during 5-year intervals.

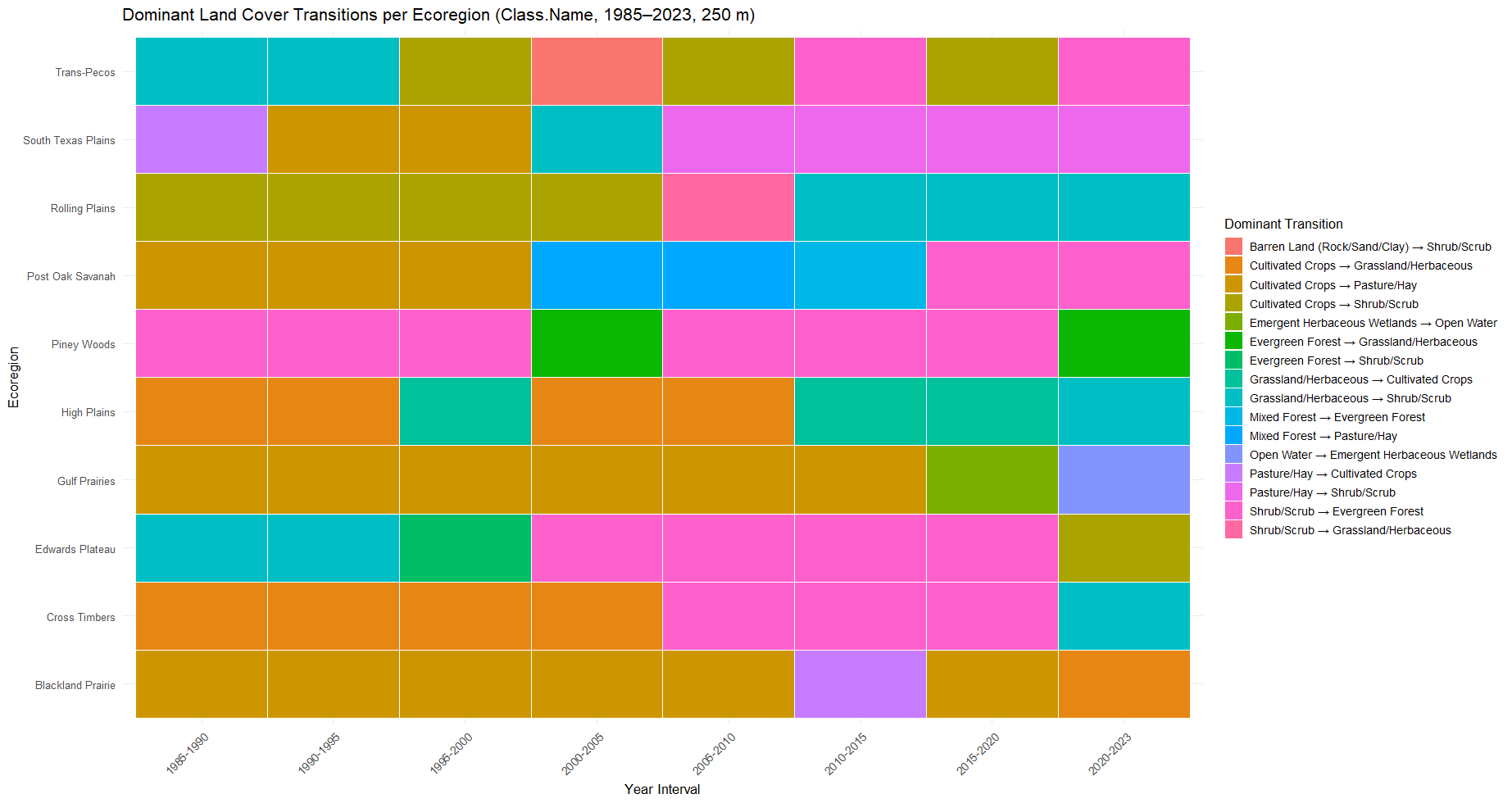

**Figure S10.** Dominant land cover transitions in each ecoregion of Texas from 1985-2023. The heatmap summarizes the most frequent land cover transition observed within each ecoregion during 5-year intervals.

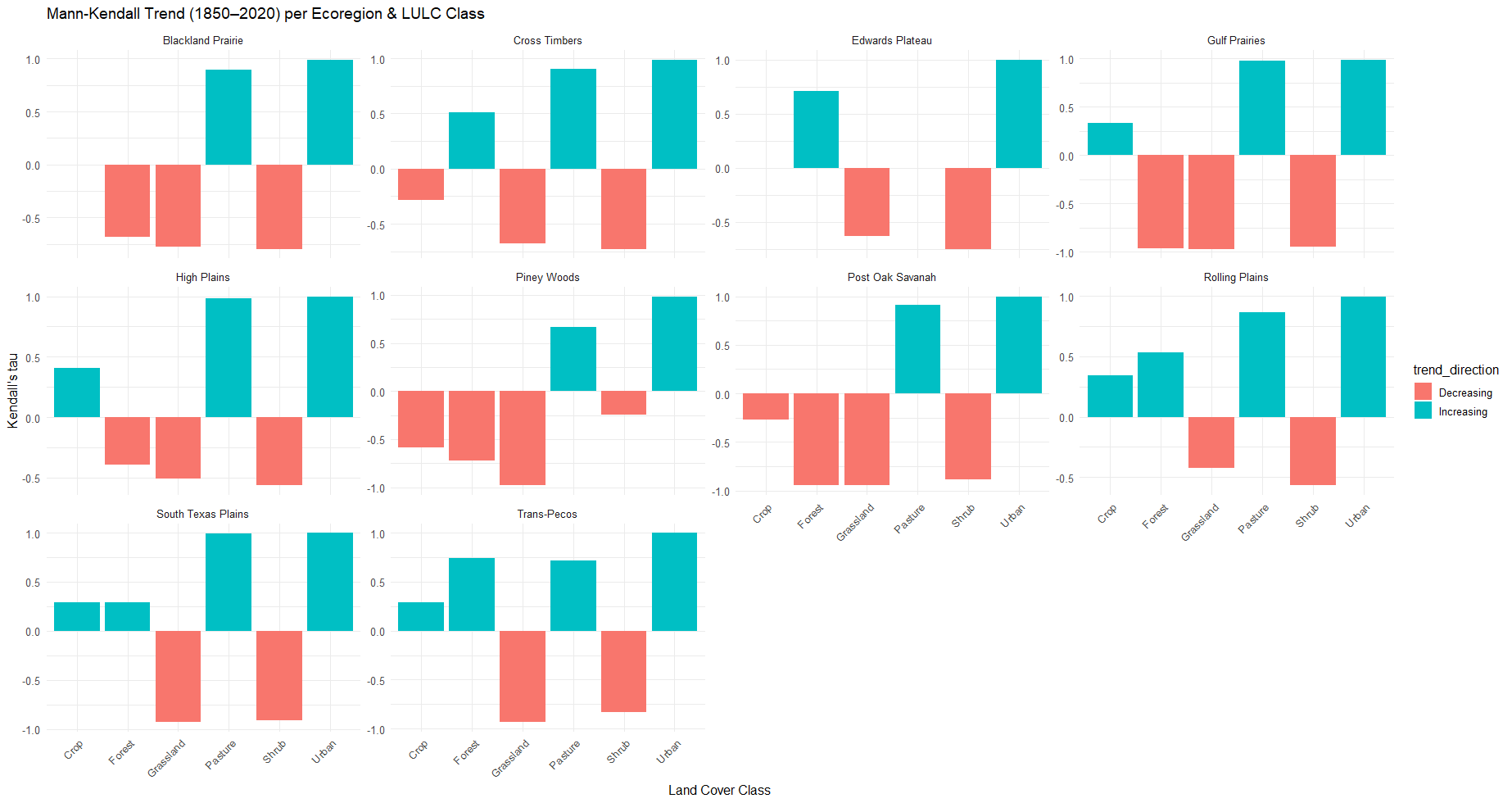

**Figure S11.** Trends in land cover classes by Texas ecoregion from 1850-2020 based on Kendall’s τ value. Each panel represents an ecoregion while bars show the direction and strength of trends. Positive values (blue) indicate trends of increase while negative values (red) indicate trends of decrease, and values near zero indicate no trend. Kendall’s τ is used to assess the trends in land cover change by determining the correlation between a habitat type and the ecoregion it is in within the time-period. The more positive the value is, the more strongly the habitat has increased its correlation with that ecoregion over time, while more negative values indicate stronger declines.

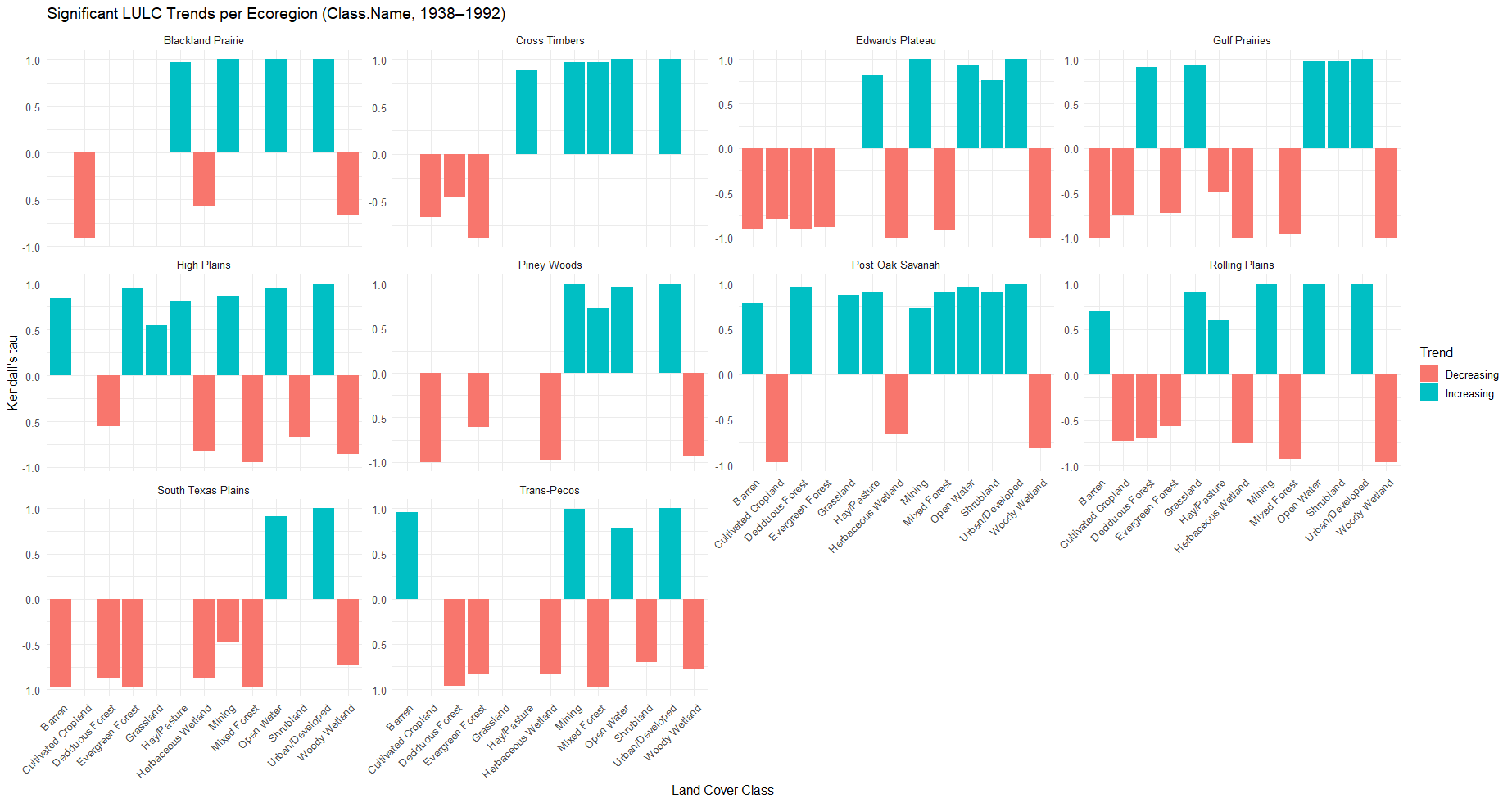

**Figure S12.** Trends in land cover classes by Texas ecoregion from 1938-1992 based on Kendall’s τ value. Each panel represents an ecoregion while bars show the direction and strength of trends. Positive values (blue) indicate trends of increase while negative values (red) indicate trends of decrease, and values near zero indicate no trend.

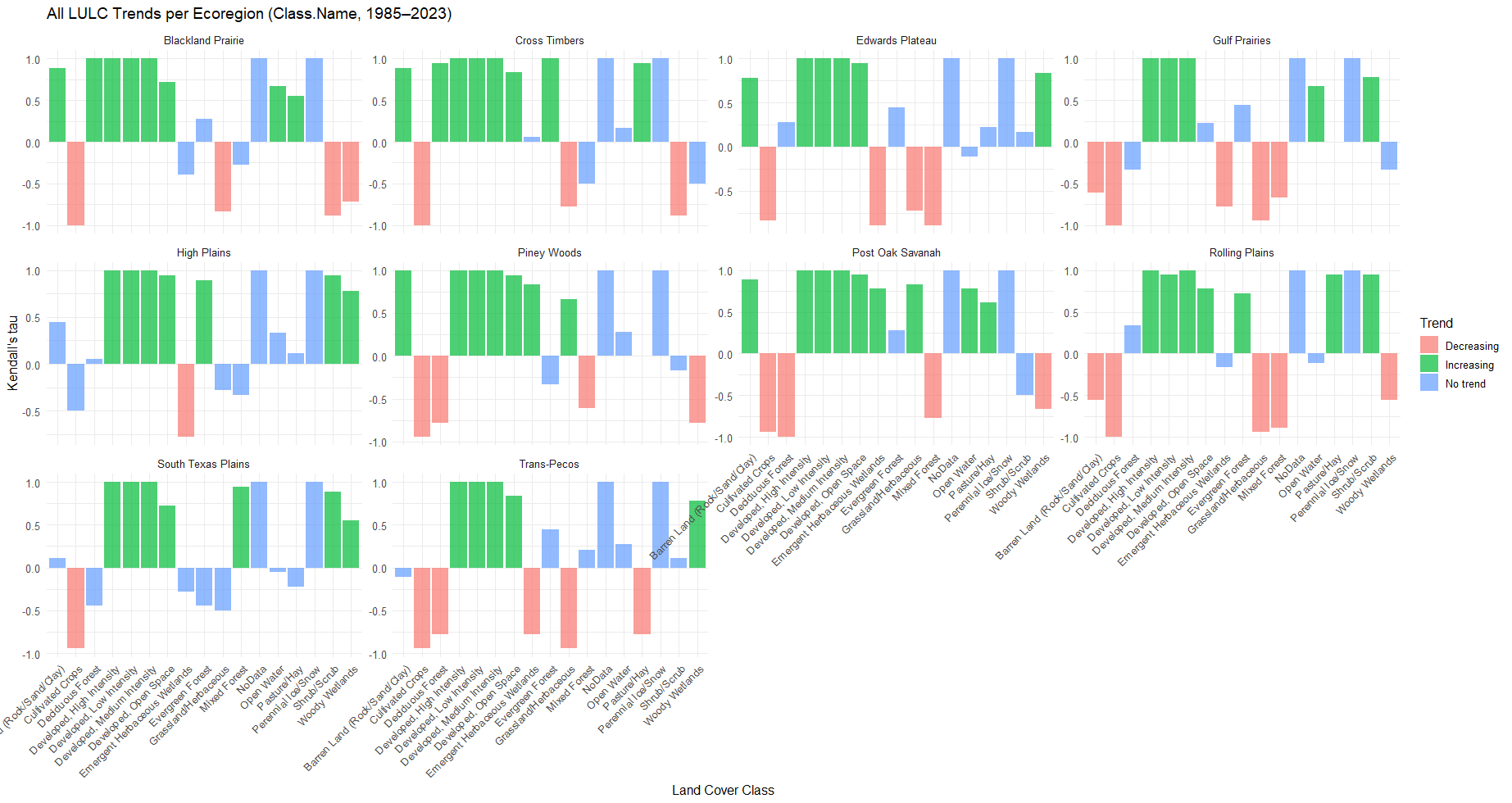

**Figure S13.** Trends in land cover classes by Texas ecoregion from 1985-2023 based on Kendall’s τ value. Each panel represents an ecoregion while bars show the direction and strength of trends. Positive values are noted in green while negative values are red and no trend is blue.

| Source | Year Period | Spatial Resolution | Number of Classes | Class List |
| --- | --- | --- | --- | --- |
| Li et al., 2023 | 1850-2020 | 1 km | 10 | No Data  Urban  Crop  Pasture  Forest  Shrub  Grassland  Wetland  Water  Barren |
| Sohl et al., 2018 | 1938-1992 | 250 m | 14 | Open Water  Urban/Developed  Mining  Barren  Deciduous Forest  Evergreen Forest  Mixed Forest  Grassland  Shrubland  Cultivated Cropland  Hay/Pasture  Herbaceous Wetland  Woody Wetland  Perennial Ice/Snow |
| U.S. Geological Survey, 2024 | 1985-2023 | 30 m | 16 | Open Water  Perennial Ice/Snow  Developed, Open Space  Developed, Low Intensity  Developed, Medium Intensity  Developed, High Intensity  Barren Land (Rock/Sand/Clay)  Deciduous Forest  Evergreen Forest  Mixed Forest  Shrub/Scrub  Grassland/Herbaceous  Pasture/Hay  Cultivated Crops  Woody Wetlands  Emergent Herbaceous Wetlands |

**Table S1.** Summary of land cover datasets used in the historical analyses across Texas. The table lists the data source (by author, year format), time range by years, spatial resolution, number of land cover classes, and class names for each dataset. The dataset by Li et al. (2023) provides a long-term modeled view of land cover from 1850-2020 at 1 km resolution with 10 generalized classes that we use in our study. The dataset by Sohl et al. (2018) covers 1938-1992 at 250 m resolution with 14 land cover classes. The dataset by the U.S. Geological Survey (2024) represents contemporary land cover from 1985-2024 at 30 m resolution with 16 classes at much higher detail.

| Common_Name | Taxa_type | Conservation_Status.IUCN | Conservation_Status.TPWD | Primary_Ecoregion | Primary_Habitat | Distribution_Range_Change |
| --- | --- | --- | --- | --- | --- | --- |
| Texas Kangaroo Rat | Mammalia | Vulnerable | Threatened | Crosstimbers | Grass | Decrease |
| Prairie Skink | Reptilia | Least Concern | Not Listed | Crosstimbers/Blackland Prairie | Grass | Decrease |
| Golden-cheeked Warbler | Aves | Endangered | Endangered | Edwards Plateau | Forest | Decrease |
| Cliff Chirping Frog | Amphibia | Least Concern | Not Listed | Edwards Plateau | Forest | Increase |
| Texas Toad | Amphibia | Least Concern | Not Listed | Generalist | Grass | Increase |
| Green Toad | Amphibia | Least Concern | Not Listed | Generalist | Shrub | Decrease |
| Eastern Tiger Salamander | Amphibia | Least Concern | Not Listed | Generalist | Wetland | Decrease |
| Western Tiger Salamander | Amphibia | Least Concern | Not Listed | Generalist | Wetland | Decrease |
| Horned Lark | Aves | Least Concern | Not Listed | Generalist | Barren | Decrease |
| Barred Owl | Aves | Least Concern | Not Listed | Generalist | Forest | Decrease |
| Blue Jay | Aves | Least Concern | Not Listed | Generalist | Forest | Increase |
| Carolina Chickadee | Aves | Least Concern | Not Listed | Generalist | Forest | Decrease |
| Eastern Kingbird | Aves | Least Concern | Not Listed | Generalist | Forest | Decrease |
| Ruby-throated Hummingbird | Aves | Least Concern | Not Listed | Generalist | Forest | Increase |
| Summer Tanager | Aves | Least Concern | Not Listed | Generalist | Forest | Decrease |
| Yellow-billed Cuckoo | Aves | Least Concern | Not Listed | Generalist | Forest | Decrease |
| American Crow | Aves | Least Concern | Not Listed | Generalist | Generalist | Increase |
| Common Grackle | Aves | Least Concern | Not Listed | Generalist | Generalist | Increase |
| Northern Mockingbird | Aves | Least Concern | Not Listed | Generalist | Generalist | Increase |
| Wild Turkey | Aves | Least Concern | Not Listed | Generalist | Generalist | Increase |
| Grasshopper Sparrow | Aves | Least Concern | Not Listed | Generalist | Grass | Decrease |
| Common Ground Dove | Aves | Least Concern | Not Listed | Generalist | Shrub | Increase |
| Greater Roadrunner | Aves | Least Concern | Not Listed | Generalist | Shrub | Increase |
| Common Gray Fox | Mammalia | Least Concern | Not Listed | Generalist | Forest | Increase |
| Eastern Fox Squirrel | Mammalia | Least Concern | Not Listed | Generalist | Forest | Increase |
| Virginia Opossum | Mammalia | Least Concern | Not Listed | Generalist | Forest | Increase |
| White-tailed Deer | Mammalia | Least Concern | Not Listed | Generalist | Forest | Increase |
| Hog-nosed Skunk | Mammalia | Least Concern | Not Listed | Generalist | Generalist | Decrease |
| Nine-Banded Armadillo | Mammalia | Least Concern | Not Listed | Generalist | Generalist | Increase |
| North American Porcupine | Mammalia | Least Concern | Not Listed | Generalist | Generalist | Increase |
| Northern Pygmy Mouse | Mammalia | Least Concern | Not Listed | Generalist | Generalist | Increase |
| Northern Raccoon | Mammalia | Least Concern | Not Listed | Generalist | Generalist | Increase |
| American Badger | Mammalia | Least Concern | Not Listed | Generalist | Grass | Range Shift |
| Eastern Mole | Mammalia | Least Concern | Not Listed | Generalist | Grass | Decrease |
| Least Shrew | Mammalia | Least Concern | Not Listed | Generalist | Grass | Increase |
| Plains Harvest Mouse | Mammalia | Least Concern | Not Listed | Generalist | Grass | Increase |
| Collared Peccary | Mammalia | Least Concern | Not Listed | Generalist | Shrub | Decrease |
| Common Muskrat | Mammalia | Least Concern | Not Listed | Generalist | Wetland | Range Shift |
| North American Beaver | Mammalia | Least Concern | Not Listed | Generalist | Wetland | Increase |
| Swamp Rabbit | Mammalia | Least Concern | Not Listed | Generalist | Wetland | Decrease |
| Green Anole | Reptilia | Least Concern | Not Listed | Generalist | Forest | Decrease |
| Southern Copperhead | Reptilia | Least Concern | Not Listed | Generalist | Forest | Decrease |
| Speckled Kingsnake | Reptilia | Least Concern | Not Listed | Generalist | Generalist | Decrease |
| Western Massasauga | Reptilia | Least Concern | Not Listed | Generalist | Grass | Decrease |
| Common Glossy Snake | Reptilia | Least Concern | Not Listed | Generalist | Shrub | Increase |
| Western Cottonmouth | Reptilia | Least Concern | Not Listed | Generalist | Wetland | Decrease |
| Killdeer | Aves | Near Threatened | Not Listed | Generalist | Generalist | Increase |
| Northern Bobwhite | Aves | Near Threatened | Not Listed | Generalist | Generalist | Decrease |
| Ornate Box Turtle | Reptilia | Not listed | Not listed | Generalist | Grass | Decrease |
| Woodhouse's Toad | Amphibia | Least Concern | Threatened | Generalist | Generalist | Increase |
| Timber Rattlesnake | Reptilia | Least Concern | Threatened | Generalist | Forest | Decrease |
| Texas Horned Lizard | Reptilia | Least Concern | Threatened | Generalist | Shrub | Decrease |
| Southern Crawfish Frog | Amphibia | Least Concern | Not Listed | Gulf Prairies | Grass | Decrease |
| Brown Pelican | Aves | Least Concern | Not Listed | Gulf Prairies | Wetland | Decrease |
| American Alligator | Reptilia | Least Concern | Not Listed | Gulf Prairies | Wetland | Decrease |
| Reddish Egret | Aves | Near Threatened | Threatened | Gulf Prairies | Wetland | Decrease |
| Black-spotted Newt | Amphibia | Vulnerable | Threatened | Gulf Prairies | Wetland | Decrease |
| White-faced Ibis | Aves | Least Concern | Not Listed | Gulf Prairies/High Plains | Wetland | Decrease |
| Green Tree Frog | Amphibia | Least Concern | Not Listed | Gulf Prairies/Piney Woods | Wetland | Decrease |
| American Mink | Mammalia | Least Concern | Not Listed | Gulf Prairies/Piney Woods | Wetland | Decrease |
| White-tailed Hawk | Aves | Least Concern | Not Listed | Gulf Prairies/South Texas Plains | Generalist | Decrease |
| Lesser Prairie Chicken | Aves | Vulnerable | Endangered | High Plains | Grass | Decrease |
| Lesser Earless Lizard | Reptilia | Least Concern | Not Listed | High Plains | Shrub | Decrease |
| Thirteen-lined Ground Squirrel | Mammalia | Least Concern | Not Listed | High Plains/Crosstimbers | Grass | Decrease |
| Red-cockaded Woodpecker | Aves | Near Threatened | Endangered | Piney Woods | Forest | Decrease |
| Marbled Salamander | Amphibia | Least Concern | Not Listed | Piney Woods | Forest | Decrease |
| Mole Salamander | Amphibia | Least Concern | Not Listed | Piney Woods | Forest | Decrease |
| Smallmouth Salamander | Amphibia | Least Concern | Not Listed | Piney Woods | Forest | Decrease |
| Spotted Salamander | Amphibia | Least Concern | Not Listed | Piney Woods | Forest | Decrease |
| Spring peeper | Amphibia | Least Concern | Not Listed | Piney Woods | Forest | Decrease |
| Woodland Vole | Mammalia | Least Concern | Not Listed | Piney Woods | Forest | Decrease |
| Northern River Otter | Mammalia | Least Concern | Not Listed | Piney Woods | Wetland | Range Shift |
| Eastern Newt | Amphibia | Least Concern | Not Listed | Piney Woods | Wetland | Decrease |
| Louisiana pine snake | Reptilia | Endangered | Threatened | Piney Woods | Forest | Decrease |
| Bachman's Sparrow | Aves | Near Threatened | Threatened | Piney Woods | Forest | Decrease |
| Black Bear | Mammalia | Least Concern | Threatened | Piney Woods/Trans-Pecos | Generalist | Increase |
| Houston Toad | Amphibia | Critically Endangered | Endangered | Post Oak Savanah | Forest | Decrease |
| Ocelot | Mammalia | Least Concern | Endangered | South Texas Plains | Shrub | Decrease |
| Mexican Chirping Frog | Amphibia | Least Concern | Not Listed | South Texas Plains | Forest | Increase |
| Texas Pocket Gopher | Mammalia | Least Concern | Not Listed | South Texas Plains | Shrub | Decrease |
| Keeled Earless Lizard | Reptilia | Least Concern | Not Listed | South Texas Plains | Shrub | Decrease |
| Mexican Tree Frog | Amphibia | Least Concern | Threatened | South Texas Plains | Forest | Decrease |
| Tropical Parula | Aves | Least Concern | Threatened | South Texas Plains | Forest | Increase |
| Texas Tortoise | Reptilia | Least Concern | Threatened | South Texas Plains | Shrub | Decrease |
| Reticulate collared lizard | Reptilia | Vulnerable | Threatened | South Texas Plains | Shrub | Decrease |
| Mexican Long-nosed Bat | Mammalia | Endangered | Endangered | Trans-Pecos | Barren | Decrease |
| Dunes Sagebrush Lizard | Reptilia | Endangered | Endangered | Trans-Pecos | Barren | Decrease |
| Canyon Tree Frog | Amphibia | Least Concern | Not Listed | Trans-Pecos | Shrub | Decrease |
| Scaled Quail | Aves | Least Concern | Not Listed | Trans-Pecos | Grass | Decrease |
| Common Black-hawk | Aves | Least Concern | Not Listed | Trans-Pecos | Wetland | Decrease |
| Round-tailed Horned Lizard | Reptilia | Least Concern | Not Listed | Trans-Pecos | Barren | Decrease |
| Long-nosed Leopard Lizard | Reptilia | Least Concern | Not Listed | Trans-Pecos | Shrub | Decrease |
| Mountain Plover | Aves | Near Threatened | Not Listed | Trans-Pecos | Barren | Decrease |
| Elf Owl | Aves | Not listed | Not Listed | Trans-Pecos | Forest | Increase |
| Spotted Bat | Mammalia | Least Concern | Threatened | Trans-Pecos | Barren | Decrease |
| Ornate Tree Lizard | Reptilia | Least Concern | Not Listed | Trans-Pecos/Edwards Plateau | Generalist | Decrease |
| Black-tailed Prairie Dog | Mammalia | Least Concern | Not Listed | Trans-Pecos/High Plains | Grass | Decrease |
| Pronghorn | Mammalia | Least Concern | Not Listed | Trans-Pecos/High Plains | Generalist | Decrease |
| Mule Deer | Mammalia | Least Concern | Not Listed | Trans-Pecos/High Plains | Generalist | Range Shift |
| Great Plains Toad | Amphibia | Least Concern | Not Listed | Trans-Pecos/High Plains | Generalist | Decrease |

**Table S2.** List of 100 Texas vertebrate species used in the historical analyses. The table provides common names and scientific names for each species; taxa type (Amphibia, Aves, Mammalia, Reptilia); conservation status (IUCN and Texas Parks and Wildlife Department); primary ecoregion; primary habitat; and observed distribution shift.

| Variable1 | Varibale2 | P-Value | Cramer’s V |
| --- | --- | --- | --- |
| Conservation_Status.IUCN | Primary_Ecoregion | 0.00 | 0.5772166 |
| Primary_Ecoregion | Primary_Habitat | 0.00 | 0.5599059 |
| Conservation_Status.IUCN | Conservation_Status.TPWD | 0.00 | 0.5474014 |
| Conservation_Status.TPWD | Primary_Ecoregion | 0.03 | 0.4658358 |
| Primary_Ecoregion | Taxa_type | 0.29 | 0.4198465 |
| Distribution_Range_Change | Primary_Ecoregion | 0.41 | 0.4070947 |
| Conservation_Status.IUCN | Taxa_type | 0.02 | 0.3486206 |
| Distribution_Range_Change | Primary_Habitat | 0.02 | 0.323059 |
| Distribution_Range_Change | Taxa_type | 0.00 | 0.3208445 |
| Conservation_Status.IUCN | Primary_Habitat | 0.18 | 0.2908835 |
| Primary_Habitat | Taxa_type | 0.12 | 0.2677203 |
| Conservation_Status.TPWD | Primary_Habitat | 0.29 | 0.2394373 |
| Conservation_Status.IUCN | Distribution_Range_Change | 0.92 | 0.1921804 |
| Conservation_Status.TPWD | Taxa_type | 0.63 | 0.1811858 |
| Conservation_Status.TPWD | Distribution_Range_Change | 0.60 | 0.1792767 |

**Table S3.** Cramér’s V association values between the different variable pairs.
